# Recombination and retroprocessing in broomrapes reveal a universal roadmap for mitochondrial evolution in heterotrophic plants

**DOI:** 10.1101/2025.02.14.637881

**Authors:** Liming Cai, Justin C Havird, Robert K Jansen

## Abstract

The altered life history strategies of heterotrophic organisms often leave a profound genetic footprint on energy metabolism related functions. In parasitic plants, the reliance on host-derived nutrients and loss of photosynthesis in holoparasites have led to highly degraded to absent plastid genomes, but its impact on mitochondrial genome (mitogenome) evolution has remained controversial. By examining mitogenomes from 45 Orobanchaceae species including three independent transitions to holoparasitism and key evolutionary intermediates, we identified measurable and predictable genetic alterations in genomic shuffling, RNA editing, and intracellular (IGT) and horizontal gene transfer (HGT) en route to a nonphotosynthetic lifestyle. In-depth comparative analyses revealed DNA recombination and repair processes, especially RNA-mediated retroprocessing, as significant drivers for genome structure evolution. In particular, we identified a novel RNA-mediated IGT and HGT mechanism, which has not been demonstrated in cross-species and inter-organelle transfers. Based on this, we propose a generalized dosage effect mechanism to explain the biased transferability of plastid DNA to mitochondria across green plants, especially in heterotrophic lineages like parasites and mycoheterotrophs. Evolutionary rates scaled with these genomic changes, but the direction and strength of selection varied substantially among genes and clades, resulting in high contingency in mitochondrial genome evolution. Finally, we describe a universal roadmap for mitochondrial evolution in heterotrophic plants where increased recombination and repair activities, rather than relaxed selection alone, lead to differentiated genome structure compared to free-living species.

## INTRODUCTION

Two critical symbiotic events define the cellular structure and mode of genetic inheritance of modern plants. The resulting organelles of these symbioses—plastids and mitochondria—marked an evolutionary revolution and established plants’ role as the primary suppliers of oxygen and organic matter on Earth. Over 1.4 billion years of evolution, the function and genetics of plastids and mitochondria have coevolved with the nucleus to perform autotrophic functions. Yet such long-term equilibrium can be broken during the evolution of parasitism. The most extreme case in eukaryotes involves the loss of mitochondrial genomes (mitogenomes) in unicellular parasites [1,2] and even the organelle itself in the microbial symbiont *Monocercomonoides* [3]. In plants, parasitism also imposes a strong selective force on energy metabolism. Parasitic plants tap into their host plants for water, nutrients, and photosynthetic products through direct haustorial connections. Their plastid genomes (plastomes) thus have elevated evolutionary rates, extensive gene loss, altered genome structure, and impaired to completely absent photosynthesis [4–7]. In contrast, few studies have attempted to address these questions in mitochondria and even fewer have identified significant differences compared to free-living species [8–10].

How mitochondrial function and genome evolution respond to heterotrophy in plants is polarized. On one hand, mitochondria play a housekeeping role in programmed cell apoptosis, calcium signaling, and oxidative homeostasis, making them fundamental to the survival and functioning of eukaryotic organisms [10]. Comparative studies in multiple parasitic and mycoheterotrophic lineages revealed minimum to no significant differences in mitochondrion-encoded gene composition compared to typical autotrophic plants [11,12]. On the other hand, there are three lines of evidence supporting potentially modified mitochondria in parasitic plants. First, theoretical models predict faster evolution in parasites in response to host-parasite arms race [13]. This was supported by higher substitution rates in parasitic plants in all three cellular genomes although limited genes were sampled [14]. Second, mitochondrial function relies on intricate genetic and biochemical crosstalk with the plastid and nucleus [5,8–10][15–17]. Thus, mitochondrial function may be influenced by the cascading effect of massive gene losses of the plastid and nuclear genomes in parasitic plants [18]. Our recent work in the parasitic Orobanchaceae revealed widespread losses of nuclear-encoded mitochondrial targeted genes in the oxidative phosphorylation (OXPHOS) pathway, which may have led to altered use of OXPHOS enzymes for mitochondrial respiration [19]. Similarly, the plastid generates numerous metabolites such as glutamate and carbohydrates to support mitochondrial respiration [17]. Substantially altered plastids thus may induce corresponding changes in mitochondria. Third, although mitochondrial function is highly conserved, several metabolic pathways are functionally redundant with forked alternative routes in plants such as the alternative oxidase [20]. This allows for more subtle and diverse outcomes of long-term relaxed selection, including not only gene losses or elevated evolutionary rates but also shifted preference for alternative metabolic pathways [21,22].

Scrutinizing signs of relaxed selection has been a central focus of molecular evolution and can be conducted by examining nucleotide substitutions in protein coding genes. Compared to the streamlined plastid, mitochondrial genes harbor a greater breadth of eccentricities such as scrambled coding regions, nonstandard genetic codes, and posttranscriptional RNA editing [23]. However, the content and evolutionary rates of core mitochondrial genes are highly conserved across most land plants. These 24 core genes comprise five ATPase genes, nine NADPH dehydrogenase genes, four cytochrome c genes, three cytochrome c oxidase genes, one membrane transport protein gene, one mature enzyme gene, and one panthenol-cytochrome c reductase gene. Nearly all parasites examined so far displayed a complete set of core genes [24–26]. However, the European mistletoe *Viscum* (Santalaceae) showed surprising losses in their NADPH dehydrogenase and ATPase genes, and highly diverged sequences for the other mitochondrial genes [9,27]. In addition to genic regions, excessive noncoding sequences are also indicative of ineffective or relaxed selection because they increase the probability of disrupting gene expression according to the mutation burden hypothesis [28]. This is manifested by increased structural rearrangements, horizontal gene transfers (HGTs), and expansion of repeats often seen in the mitogenomes of parasitic plants [11,12,25,29–31]. These macro- and microstructural genomic modifications thus often covary with evolutionary rates and selection pressure, which is well established in the plastomes of parasitic plants (e.g., Wicke et al., 2016) but remain to be tested in mitogenomes.

Investigation on mitogenome evolution in parasitic plants has largely focused on lineages with a single origin of parasitism and has been restricted to coding regions [11,31]. Challenges in mitogenome assembly and annotation further hindered broader taxon sampling to identify key points of selective transition in relation to shifts in life history strategies [32]. Here, we focused on the broomrapes (Orobanchaceae), a group of cosmopolitan and diverse parasites in Lamiales. Members of this family span the entire spectrum of plant parasitism from free-living species to chlorophyllous hemiparasites and achlorophyllous holoparasites devoid of photosynthesis capacity. The three independent origins of holoparasitism in Orobanchaceae provide a suitable comparative framework to explore the relationship between shifts in life history strategy and mitogenome evolution. Furthermore, cryptic hemiparasites such as *Harveya* and *Lathraea* are evolutionary intermediates that perform photosynthesis only at certain life stages, which can reveal the precise evolutionary trajectory of mitogenomes to a fully heterotrophic lifestyle. Therefore, we used comparative genomic techniques to explore the adaptive and nonadaptive processes shaping nucleotide substitution and mitogenome structure in Orobanchaceae.

## RESULTS

### Discordant organellar phylogenies provide insights into Orobanchaceae divergence

We assembled and annotated mitogenomes and plastomes from 45 Orobanchaceae species representing 28 genera (Table S1). Both the mitochondrial and plastid phylogeny supported three independent origins of holoparasitism (Fig. 1). Among these three lineages, tribe Orobancheae *sensu* McNeal et al. [33], consisting of *Orobanche, Aphyllon, Phelipanche, Cistanche, Mannagettaea, Conopholis, Epifagus,* and *Kopsiopsis*, was placed as sister to tribe Buchnereae with moderate support (79 and 87 ultrafast bootstrap in IQ-TREE, referred to as UFBP hereafter). The other two holoparasitic lineages were well nested within tribes Buchnereae (*Aeginetia, Hyobanche, Christisonia*, and *Harveya*) and Rhinantheae (*Lathraea*) with maximum support. Several major conflicts existed between the mitochondrial and plastid phylogenies. For example, *Brandisia* was placed as sister to tribe Pedicularideae with 99 UFBP based on the mitochondrial phylogeny but recovered as sister to Buchnereae with 100 UFBP in the plastid phylogeny. Similarly, *Monochasma* was placed as sister to Buchnereae in the mitochondrial phylogeny (89 UFBP), but inferred to be sister to all parasitic Orobanchaceae using plastid (73 UFBP). At the species level, the two *Aphyllon fasciculatum* accessions (voucher JEPS 127839 and voucher I. W. Clokey and B. G. Anderson) formed a clade in the mitochondrial phylogeny (100 UFBP), but were paraphyletic in the plastid phylogeny with maximal support. These well-supported conflicts are indicative of a deep history of hybridization involving *Brandisia* and *Monochasma*, as well as potentially widespread host-driven cryptic speciation exemplified in *Aphyllon fasciculatum*. A detailed discussion of the phylogenetic insights of these results is provided in Supplementary Note 1.

**Figure 1.**
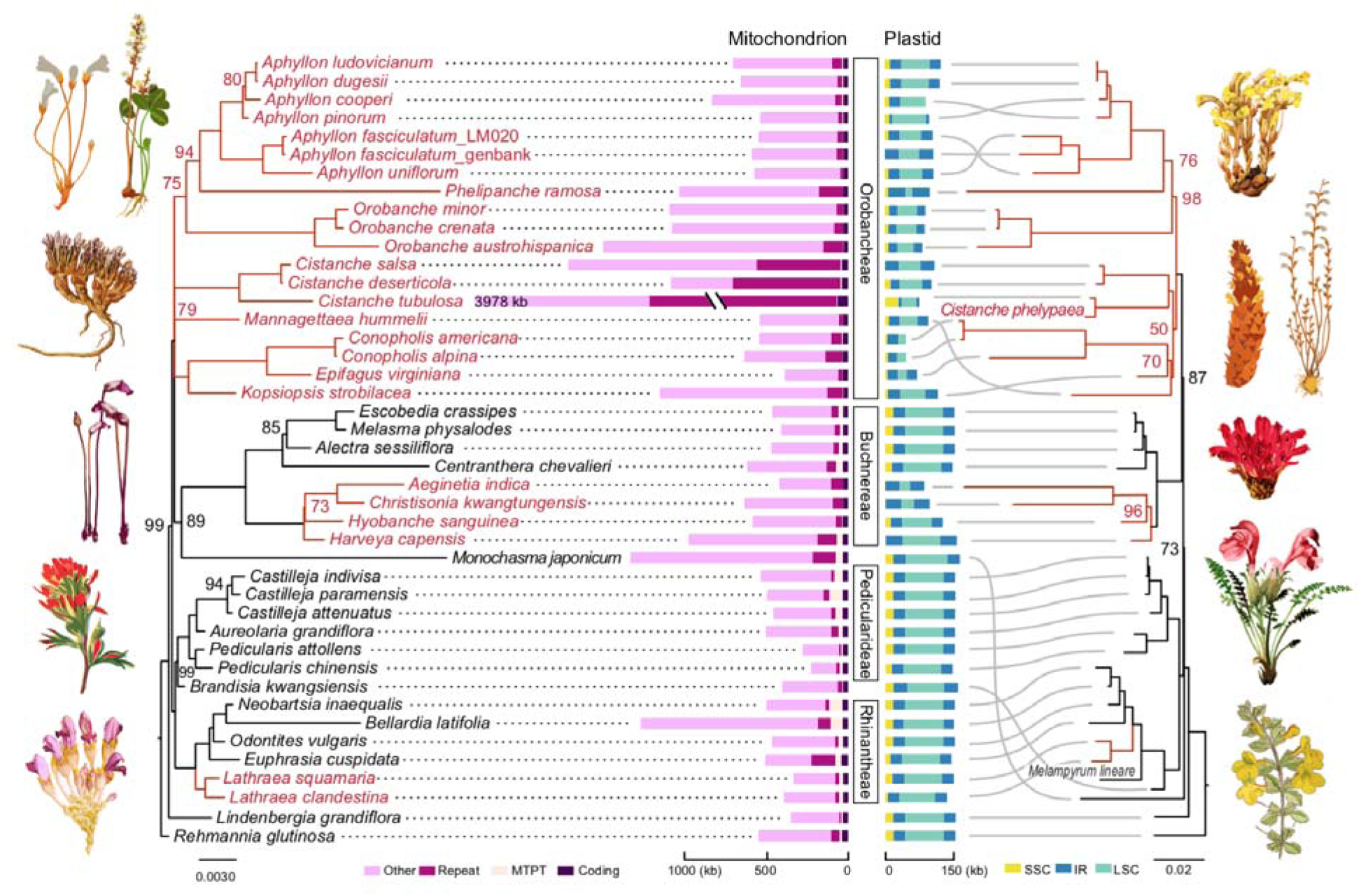
Mitochondrial and plastid genome structure in Orobanchaceae. Maximum likelihood phylogenies of Orobanchaceae were inferred from mitochondrial (left) and plastid (right) genes using IQ-TREE. Branch support was evaluated by 1000 ultrafast bootstrap replications and only support values lower than 100 are indicated at nodes. Tribe names are labeled in the middle. Phylogenetic incongruences are shown by gray lines and holoparasitic lineages are highlighted in red. Mitochondrial genomes are color-coded by sequence types including coding regions (dark purple), mitochondrial DNA of plastid origin (MTPT, salmon), repeats (purple), and others (pink). Plastid genomes are color-coded by small single-copy region (SSC, yellow), large single-copy region (LSC, aqua), and inverted repeat (IR, blue). Illustrations of plants were obtained from the public domain of the Biodiversity Heritage Library (https://www.flickr.com/photos/61021753@N02/). From top left to bottom right: *Aphyllon uniflorum*, *Orobanche minor, Mannagettaea hummelii, Aeginetia indica, Castilleja coccinea, Lathraea clandestina, Aphyllon fasciculatum, Conopholis americana, Epifagus virginiana, Hyobanche sanguinea, Pedicularis portenschlagii, and Lidenbergia grandiflora*.

### Altered genome size and gene content in the plastid but not mitochondrion

Plastomes in Orobanchaceae ranged from 45.8 kb to 164.1 kb in size (Table S2). The most compact genomes were found in non-photosynthetic lineages, which were often coupled with profound structural rearrangements such as the loss of inverted repeats in *Conopholis* (Fig. 1). Plastome size reduction was also correlated with decreased GC content, although not reaching statistical significance (phylogenetic generalized least squares PGLS *p*-value = 0.085; Fig. S1A). When assessing copy number using base coverage, holoparasitic species on average contained 217.1±235.1 plastomes per cell showing no significant difference to hemiparasites (count = 327.4±261.2; phylANOVA *p-*value = 0.786; Table S3). Other aspects of plastome degradation are largely consistent with previous investigations in Orobanchaceae [5,6,34], and will not be further discussed.

Mitogenomes in Orobanchaceae ranged from 225.6 kb in *Pedicularis* to 3.98 Mb in *Cistanche* (Table S4). The median size of the mitogenome was 547.0 kb and was relatively conserved at the genus level but variable across the family (Fig. 1). No significant size difference was observed between species displaying different modes of parasitism (i.e., holoparasite versus photosynthetic species; phylANOVA *p-*value = 0.488). Mitogenome GC content was conserved across the family (42.4–48.0%) but unlike plastome, showed no correlation with genome size (Fig. S1B), suggesting that disparate processes have shaped the nucleotide compositions of the two organelles.

The core set of 24 mitochondrial genes was intact in all species except for a pseudogenized *mttB* in the holoparasitic *Mannagettaea hummelii* (Table S5). Putative loss of function in this *mttB* copy was supported by the presence of a premature stop codon and lack of sequence homology at the 5’ end. Phylogenetic investigation on the 5’ end upstream sequence pointed to a host-derived insertion from the legume genus *Dalbergia* (Fabaceae) as the main culprit of gene dysfunction (Fig. S2A). It also pointed out two additional HGTs from Fabaceae to other independently evolved holoparasites at this locus, including *Cynomorium* (Cynomoriaceae) and *Lophophytum* (Balanophoraceae) (Fig. S2A). Read mapping strongly corroborated the presence of this premature stop codon in the mitochondrial *mttB* in *Mannagettaea* and suggested a putatively intact nuclear copy indicating transfer to the nuclear genome, but the sequencing coverage was inadequate to recover its full length (Fig. S2C).

Non-core mitochondrial genes in Orobanchaceae including ribosomal proteins and succinate dehydrogenase (*sdh3* and *sdh4*) are frequently lost or pseudogenized in the mitogenome (Table S5) and like other angiosperms, potentially transferred to the nucleus [35]. Eight members of the large (*rpl5, 10, 16*) and small (*rps3, 4, 10, 12, 14*) ribosomal subunits were conserved in most species (Table S5). For all genes, intron content was conserved within Orobanchaceae with most of the dynamics found in *cox2i691*. This cis-spliced intron has been lost at least three times and substituted by host-derived HGTs twice (Table S5). Two potential shifts from cis-to-trans splicing were found in *ccmFci829* of *Hyobanche sanguinea* and *nad5i1872* of *Pedicularis attolens* as evidenced by the long distance between adjacent exons (21.1–41.9 kb).

### Accelerated structural rearrangements in holoparasites

Interspersed repeats on average accounted for 7.0% of the mitogenome in Orobanchaceae, and there was no significant difference between the two modes of parasitism (phylANOVA *p*-value = 0.45). However, the highest proportion of repeats was commonly found in holoparasitic lineages including *Cistanche* (30.1–70.5%), *Conopholis* (12.2–16.3%), and *Aeginetia* (17.8%; Table S4). The hemiparasitic *Euphrasia* also carried disproportionately abundant repeats (29.0%) and was an outlier compared to other hemiparasites. Individual repeat units ranged from 98 bp to 21.1 kb in size with a median length of 192 bp (Table S6). Among the 451 structural repeats identified from the De Bruijn assembly graph, 300 were singletons lacking homology in other species, but 92 of the remaining repeats convergently evolved from homologous regions (Fig. S3). These 92 repeats can be clustered into 24 groups and the biggest cluster comprised 9 parallel origins of repeats from the same ca. 100 bp region that resembles *nad* gene sequences (Cluster 7 in Table S6).

To further investigate mitochondrial structural dynamics, we conducted pairwise genome alignment with comparison to the free-living species *Rehmannia glutinosa* in Orobanchaceae. Within 35 million years of evolution [36], synteny was barely detectable outside coding regions in all species (Fig. S4). Yet holoparasites in particular showed more extensive rearrangements and accelerated synteny erosion (phylANOVA *p*-value = 0.05 based on the percentage of mitogenome in synteny; Figs. S4-6). For example, a 12.7 kb syntenic segment spanning *trnS-GGA*, *trnD-GUC*, and *nad4* was almost universally conserved in hemiparasites, but this segment was truncated to 5.6-10.8 kb in holoparasites (Fig. S4). In general, whole-genome pairwise alignments demonstrated shorter and more scattered syntenic segments in holoparasites (Fig. S5). This was further corroborated by the steep curve of the cumulative length of syntenic segments in hemiparasites compared to holoparasites (Fig. S6). The smoother curves and lower end points suggested that holoparasites on average lack long syntenic segments and have fewer sequences in alignment with their ancestors.

### Retroprocessing and mutational bias attenuate RNA editing in holoparasites

An average of 385.3±9.0 cytidines to uridines (C-to-U) RNA editing sites per species were predicted bioinformatically for the 24 core mitochondrial genes (Table S7). Most of these sites were located in the second codon position (64.1%) with 29.3% and 6.6% found in the first and third codon positions, respectively. In general, holoparasites have lost more RNA editing sites (mean editing sites = 391.6) compared to hemiparasites (mean = 399.1; phylANOVA *p-*value = 0.289). Further investigation revealed that such a bias was attributable to both RNA-mediated retroprocessing and biases in point mutations (Fig. 2). First, seven out of the ten events of retroprocessing-mediated loss of RNA editing sites were associated with holoparasitic lineages (Fig. 2B). In these regions, continuous stretches of C-to-T mutations are identified in *atp6*, *cox2*, and *nad4L,* each containing up to 15 editing sites spanning 191 to 512 bp of mitochondrial DNA (Fig. S7). This mechanism was further corroborated by three repeated losses of a group I intron flanking the RNA editing sites in *cox2*, pointing to gene conversion of reverse-transcribed RNA as the molecular mechanism (Fig. 2B). The case involving *cox2* and shared by the common ancestor of *Lathraea* and *Neobartsia* suggests that these historical events can date back to their common ancestor in the Late Miocene [36]. Second, mutational biases have also contributed to variations in RNA editing. By inferring ancestral sequences and mapping C-to-T mutations to the species tree, we found that holoparasitic lineages are three times more likely to accumulate C-to-T mutations when corrected for background mutation rate (Spearman rho = 0.656 vs. 0.222 in holoparasites vs. hemiparasites; Fig. 2A).

**Figure 2.**
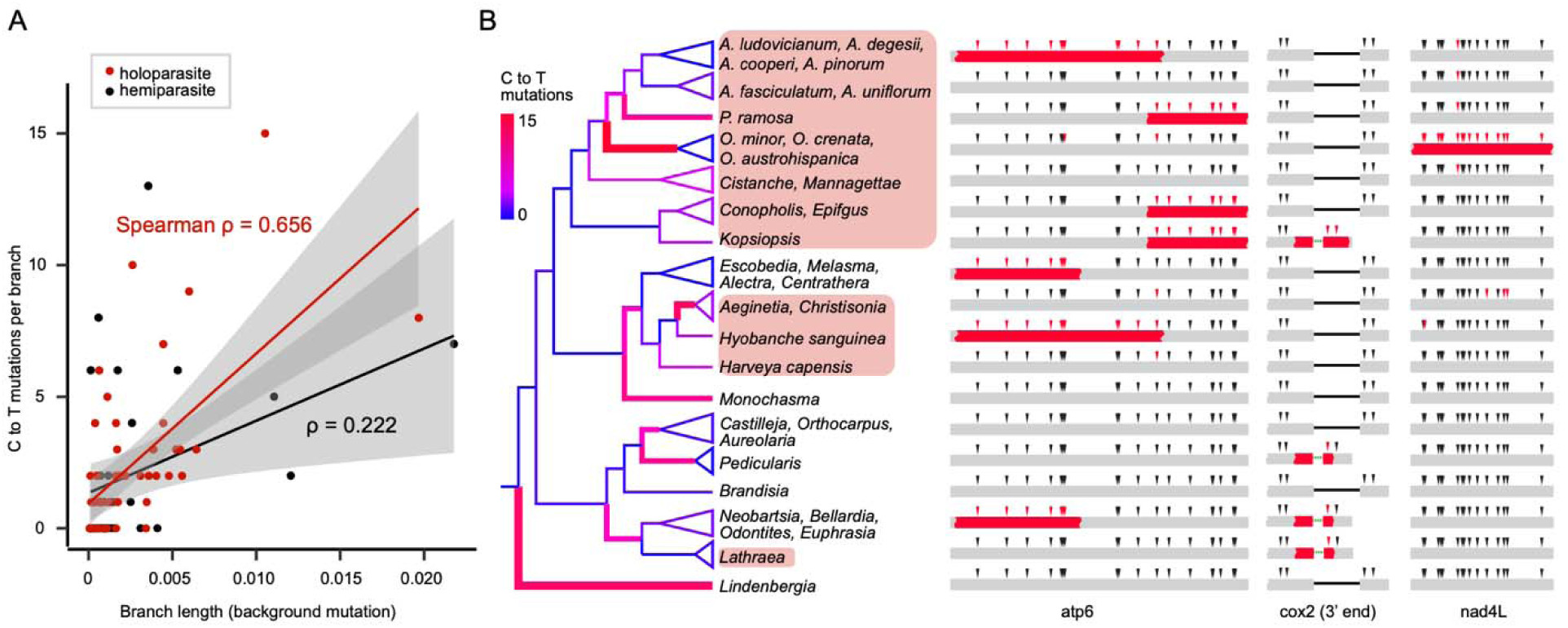
Higher rate of C-to-T point substitutions and retroprocessing lead to rapid loss of RNA editing sites in holoparasitic Orobanchaceae. (A) Number of C-to-T mutations in RNA editing sites per branch plotted against their branch length in mutation units (nucleotide substitutions per site). C-to-T mutations are inferred using ancestral state reconstruction for the internal branches and only sites not nested within a potentially retroprocessed genetic block are included. (B) Ten independent cases of retroprocessing lead to large-scale loss of RNA-editing. The phylogeny on the left is color-coded by the count of C-to-T mutations in RNA editing sites for each branch. Taxa highlighted in red are holoparasitic. Predicted RNA-editing sites are shown as triangles on each gene and the red color indicates loss of RNA-editing. Multiple continuous RNA-editing losses, as well as losses of flanking introns in *cox2*, are best explained by retroprocessing.

### Minimum intracellular gene transfer in non-photosynthetic angiosperms

Gene flow from plastids to mitochondria (MTPT) is one of the many mechanisms for mitochondria to uptake non-native DNA. These intracellular transfers accounted for a small proportion of mitogenome in Orobanchaceae (0.53–17.50%), with the lowest MTPT content found in holoparasites (Fig. 3A). Further statistical tests suggested a significant difference between holoparasitic and photosynthetic species (phylANOVA *p-*value = 0.019). When we expanded the comparison to additional clades across the Plant Tree of Life, five independently evolved non-photosynthetic lineages exhibited significantly lower MTPT content compared to their photosynthetic relatives (Fig. 3A; Table S8; phylANOVA *p-*value = 0.012). These lineages included not only holoparasites such as *Ombrophytum* (Balanophoraceae, Santalales) and *Hydnora* (Hydnoraceae, Piperales) but also mycoheterotrophs such as the ghost pipe *Monotropa* (Ericaceae, Ericales). Furthermore, cryptic Orobanchaceae hemiparasites such as *Lathraea* and *Harveya capensis*, which highly resemble leafless holoparasites but conduct photosynthesis at early life stages [37,38], exhibited high MTPT content comparable to other hemiparasites (4.19–8.85%; Fig. 3A). These results allude to universally altered interactions between plastids and mitochondria triggered by the loss of photosynthesis.

**Figure 3.**
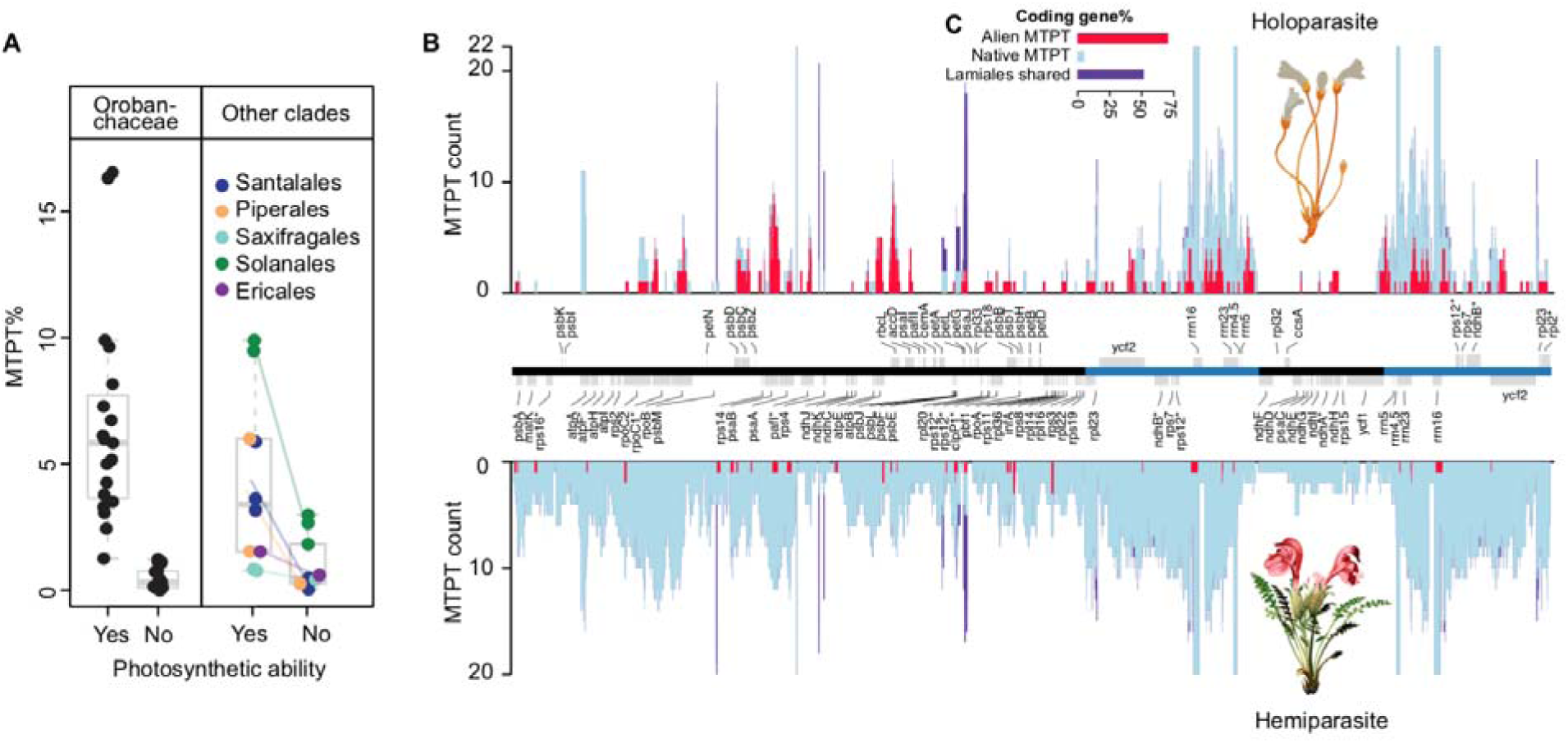
Parallel reduction of mitochondrial plastid (MTPT) content in non-photosynthetic plants caused by reduced intracellular gene transfer. (A) Reduced MTPT content in non-photosynthetic Orobanchaceae (left) and other angiosperm clades (right) including Balanophoraceae (Santalales), Hydnoraceae (Piperales), Cynomoriaceae (Saxifragales), *Cuscuta* (Convolvulaceae, Solanales), and *Monotropa* (Ericaceae, Ericales; mycoheterotrophic). A full list of species is provided in Table S8. (B) Holoparasitic Orobanchaceae receive less native MTPT but more alien MTPT. Using maximum likelihood phylogeny, MTPTs are classified into alien MTPTs from non-Lamiales species (red), native MTPTs transferred from Orobanchaceae (light blue), or ancestral MTPTs universally present in Lamiales mitochondria (purple). Homologous regions of MTPT insertions for holoparasitic (top) and hemiparasitic and free-living Orobanchaceae (bottom) are mapped to the plastome of *Rehmannia glutinosa* (middle). The inverted repeat regions are highlighted in blue. (C) Percentages of MTPTs nested within plastid coding regions. MTPTs were classified and color-coded by the same three types in (B): alien MTPT (red), native MTPT (light blue), and ancestral MTPT in Lamiales (purple).

To further investigate the patterns and mechanisms of MTPT purging in holoparasitic Orobanchaceae, we inferred the genetic background and genomic location for all 1,094 MTPTs using phylogenetic approaches. The results suggested that native plastomes on average accounted for 67.2% of the MTPT, while plastid transfers from outside Orobanchaceae contributed 17.4%. Such a proportion is sharply contrasted when partitioned by life history strategy—a minimum 6.0% of the MTPTs are from alien sources in hemiparasites, but this number rises to 28.8% in holoparasites (phylANOVA *p-*value = 1.3e-3). The comprehensive collection of plastid sequences on GenBank allows us to pinpoint the donor lineages of 48 alien MTPTs to genus level and 80 more to family level with >85 UFBP (Table S9; Data S1). These donors are strongly associated with hosts such as the insertion of *Quercus* plastid DNA in *Conopholis americana* and *Tamarix* in *Cistanche tubulosa* (Table S9). Besides direct plastid-to-mitochondrion transfers, 16.5% of MTPT are ancestral intracellular gene transfers shared broadly in Lamiales (Fig. S8D) or results from more complex host plastid to host mitochondrion to parasite mitochondrion transfer based on phylogeny (Fig. S8C; Supplementary Note 1; Data S1).

Mapping MTPTs to the plastome revealed a striking disparity in transferability across the genome (Fig. 3B). For alien MTPT, 73.7% (n = 112) of the sequences reside strictly within coding regions and are evenly distributed throughout the plastome (Fig. 3B-C). This suggests a stochastic and potentially RNA-mediated mechanism. In direct contrast, only 5.7% (n = 42) of the native MTPT sequences are within plastid genes and they demonstrate strongly biased genomic distribution—the inverted repeats are hotspots of gene transfer while the single copy regions are virtually deserts. Especially in holoparasites, the two single-copy regions account for 66.7% of the plastome but contribute only 13.9% (16.4 kb) of the total MTPTs, whereas the inverted repeats contribute 101.8 kb in total (Fig. 3B).

### HGTs inflate mitogenome sizes

In addition to plastid sequences, most non-native mitochondrial sequences come from the mitogenome of other species. Our newly developed HGTScanner program (see Methods) can identify HGT in both genic and intergenic regions and precisely pinpoint the donor lineage. Across 45 Orobanchaceae species, we discovered 6,504 high confidence HGTs. Investigation of the HGT donor lineages revealed a fascinating history of host-parasite coevolution and was discussed elsewhere (Cai *et al,* unpublished; see supplemental attached for review only and not for final publication). These alien mitochondrial sequences (excluding MTPTs) accounted for 1.8% to 29.8% of the mitogenome in each species and were positively correlated with the mitogenome size (PGLS *p*-value = 4.643e-06; Fig. S9). Within hemiparasites, HGT on average accounted for 5.2±3.2% of the mitogenome with a median segment size of 185.3 bp (Fig. S9; Table S4). This moderate level of HGTs is only slightly higher than the free-living species at 2.4–3.1%. In sharp contrast, HGTs in holoparasites expanded by more than twofold (13.5±5.4%; phylANOVA *p-*value =0.048), which is attributable to their higher abundance (218.5 per genome; Fig. S9) and longer segment size compared to those from the hemiparasites (218.0 bp; Fig. S10). In the three cryptic hemiparasites *Harveya capensis, Lathraea clandestine*, and *Lathraea squamaria,* HGT accounts for 8.7–13.4% of the mitogenome respectively, which are in close alignment with holoparasites rather than hemiparasites.

### Accelerated molecular evolution and relaxed selection in plastomes, not mitogenomes of parasites

We calculated the average synonymous (*d_S_*), nonsynonymous (*d_N_*) substitution rates, and selective pressure ω (*d_N_/d_S_*) of each species by comparing concatenated organellar genes to the free-living *Rehmannia*. A five-fold difference is found in mitochondrial *d_S_* across Orobanchaceae ranging from 0.0061 to 0.0288, which is even higher than the three-fold rate difference in their plastids (Fig. 4A). Plastid *d_S_* and ω are significantly higher in holoparasites compared to hemiparasites (phylANOVA *p-*value < 0.034), but we found no significant lifestyle-linked difference for mitochondrial *d_S_* and ω (phylANOVA *p-*value > 0.455). Species with higher plastid *d_S_* tend to have higher mitochondrial *d_S_*, although statistically trivial (PGLS *p-*value >0.16; Fig. 4A). We found no correlation between plastid ω and mitochondrion ω (PGLS *p-*value >0.79; Fig. 4B), but the hemiparasite *Monochasma japonicum* had exceptionally high *ω* in its plastid genes, which is likely a result of excessive posttranscriptional RNA editing instead of relaxed selection.

**Figure 4.**
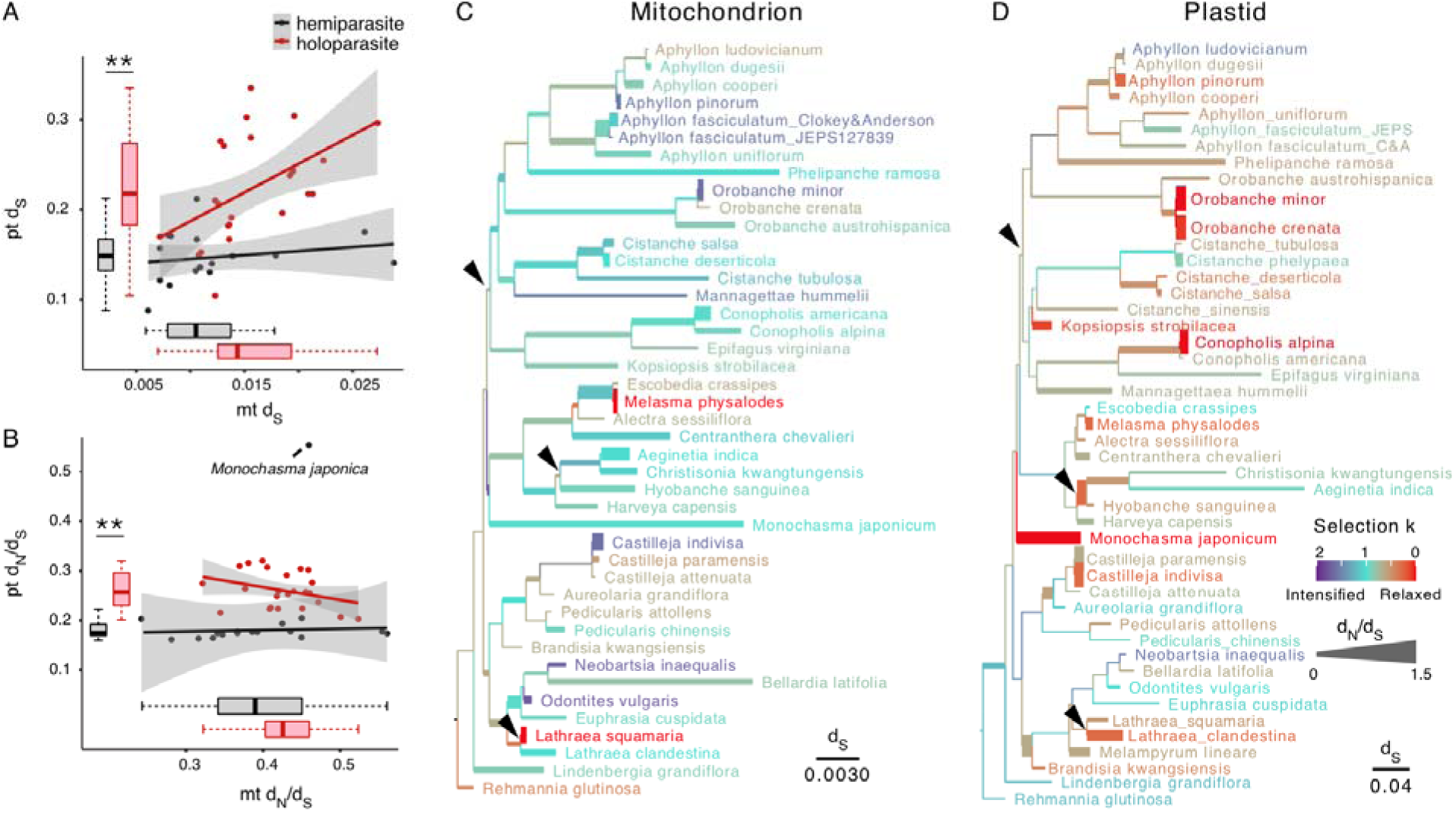
Relaxed selection in plastomes but not mitogenomes during the transition to holoparasitism. (A) Mitochondrial and plastid *d_S_* in hemi- and holo-parasites. The dot plot shows the correlation between mitochondrial and plastid *d_S_* for hemiparasites (black) and holoparasites (red). Rates are inferred from pairwise comparisons to the free-living *Rehmannia glutinosa.* The asterisks above the box plot indicate a significant difference based on phylogenetic ANOVA (*p-*value <0.05). (B) Correlation between mitochondrial and plastid *d_N_*/*d_S_*. Note the unusually high *d_N_*/*d_S_* value in the hemiparasite *Monochasma japonica*. (C–D) Shifts of selective strength for mitochondrion (C) and plastid (D). Branch length is scaled by *d_S_* and arrowheads mark three independent transitions to holoparasitism. The selection strength parameter *k* and *d_N_/d_S_* are inferred under the free-ratio model in HYPHY and CODEML, respectively. Branches are color-coded according to *k*. Low *k* (<1, red–brown) indicates relaxed selection; *k* >1 (blue-purple) suggests intensified selection. Branch widths are proportional to *d_N_/d_S_*.

To more finely characterize clade-specific selection shifts, we applied the free-ratio model to estimate ω and the selection parameter *k* defined in RELAX [39]. For plastid genes, all three holoparasitic lineages have high ω and low *k* values that are indicative of relaxed selection (thick branches with warm colors in Fig. 4D). Some hemiparasites, including *Monochasma* and *Castilleja*, also exhibit substantial relaxed selection in their plastid genes. Selection in mitochondrial genes is more subtle and most branches evolved nearly neutrally (ω and *k* close to 1; Fig. 4C). Species like *Lathraea squamaria* and *Melasma physalodes* are among the few outliers showing exceptional levels of relaxed selection in their mitochondrial genes (Fig. 4C).

To identify gene-specific selection shifts and evaluate their statistical significance, we partitioned mitochondrial genes into functional groups and compared models with alternative branch partitioning using likelihood ratio tests (LRT) in HYPHY (Table 1). Significant relaxed selection was identified in *cob, ccm*, and *mttB* in various holoparasitic clades compared to photosynthetic species (*k* = 0.63–0.89; LRT *p-*value <0.05; Table 1; Table S10). But *nad* genes encoding the NADPH dehydrogenase genes showed significantly intensified purifying and positive selection among parasitic Orobanchaceae compared to free-living species (*k* = 2.20, LRT *p*-value = 4.06e-3). The *cox* genes encoding the cytochrome c oxidase also showed significantly intensified selection in holoparasites (*k* = 1.62, LRT *p*-value = 3.04e-5). These heterogeneities in the direction of selection among different genes led to the lack of signal in the concatenated mitochondrial sequences (LRT *p-*value = 0.53).

**Table 1.**
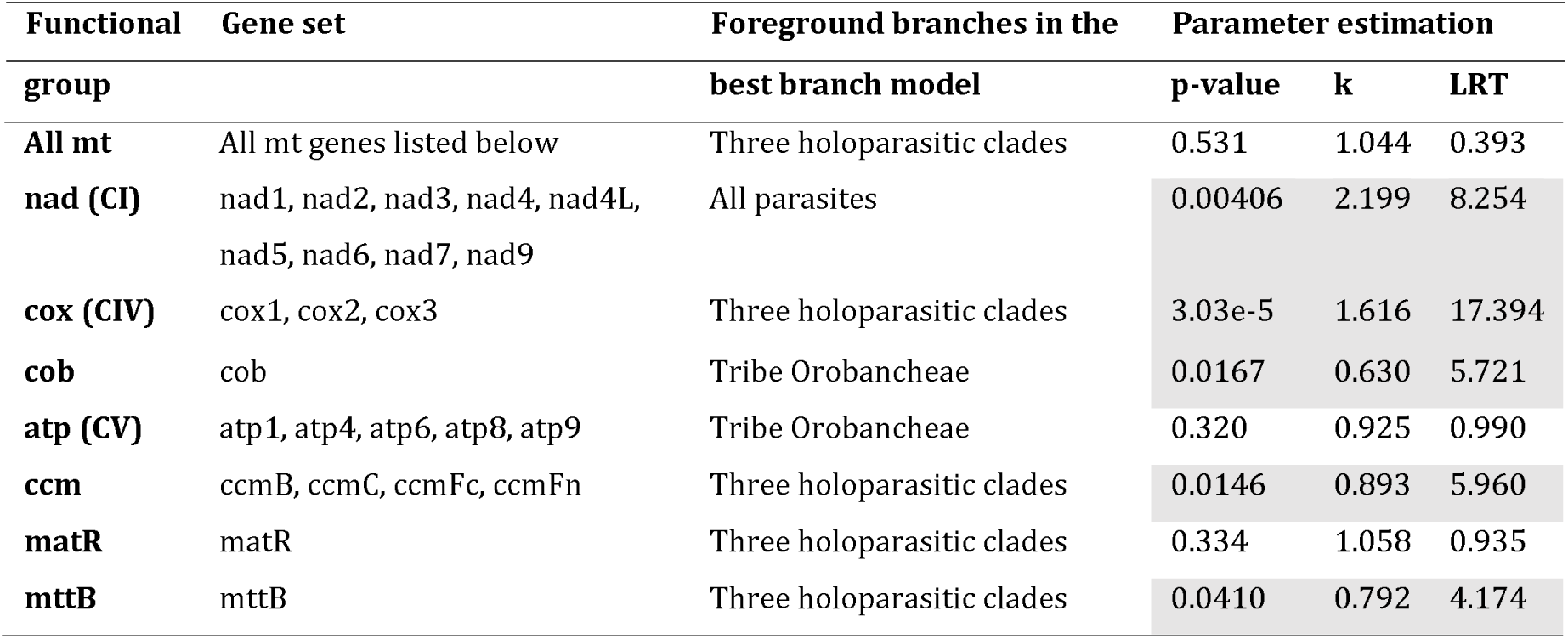
Gene-specific assessment of lifestyle effects on selection using the selection intensity parameter *k* defined in RELAX. The RELAX analyses were based on concatenated mitochondrial genes in different functional groups. Likelihood ratio tests (LRT) were performed to identify the best branch partition model and evaluate its goodness-of-fit compared to the null model assuming *k* = 1 for all branches. A significant *p*-value (<0.05) and *k* < 1 indicates relaxed selection in the foreground branches; *k* > 1 indicates intensified selection. Genes with significantly altered selection in foreground branches are highlighted in gray.

### Correlation of nucleotide substitutions and genomic traits

The correlation between mitochondrial nucleotide substitutions (*d_S_*, ω, GC%) and genomic structural changes (size, repeats, synteny, RNA editing, MTPT, HGT) was assessed under the multivariate phylogenetic comparative framework implemented in COEVOL [40]. The results indicated that none of the nucleotide substitution traits were intercorrelated (i.e., *d_S_,* ω, GC%), but many were strongly linked with genomic structural changes (Table 2). For example, mitochondrial *d_S_* was positively correlated with genome size (posterior probability of marginal correlations pp_MC_ = 0.91 and maximally controlled correlations pp_MCC_ = 0.87; Table 2). The GC content was higher in larger genomes (pp_MC_ = 0.93; pp_MCC_ = 0.75) and high HGT content (pp_MC_ = 0.94; pp_MCC_ = 0.97). Mitogenome size was strongly positively linked with MTPT, HGT, and repeat content (pp_MC_ = 0.99-1; pp_MCC_ = 0.97–0.98). Increased genomic rearrangements measured by the proportion of syntenic regions were strongly positively correlated with HGT and MTPT (pp_MC_ = 0.98-0.99; pp_MCC_ = 0.92–0.95).

**Table 2.**
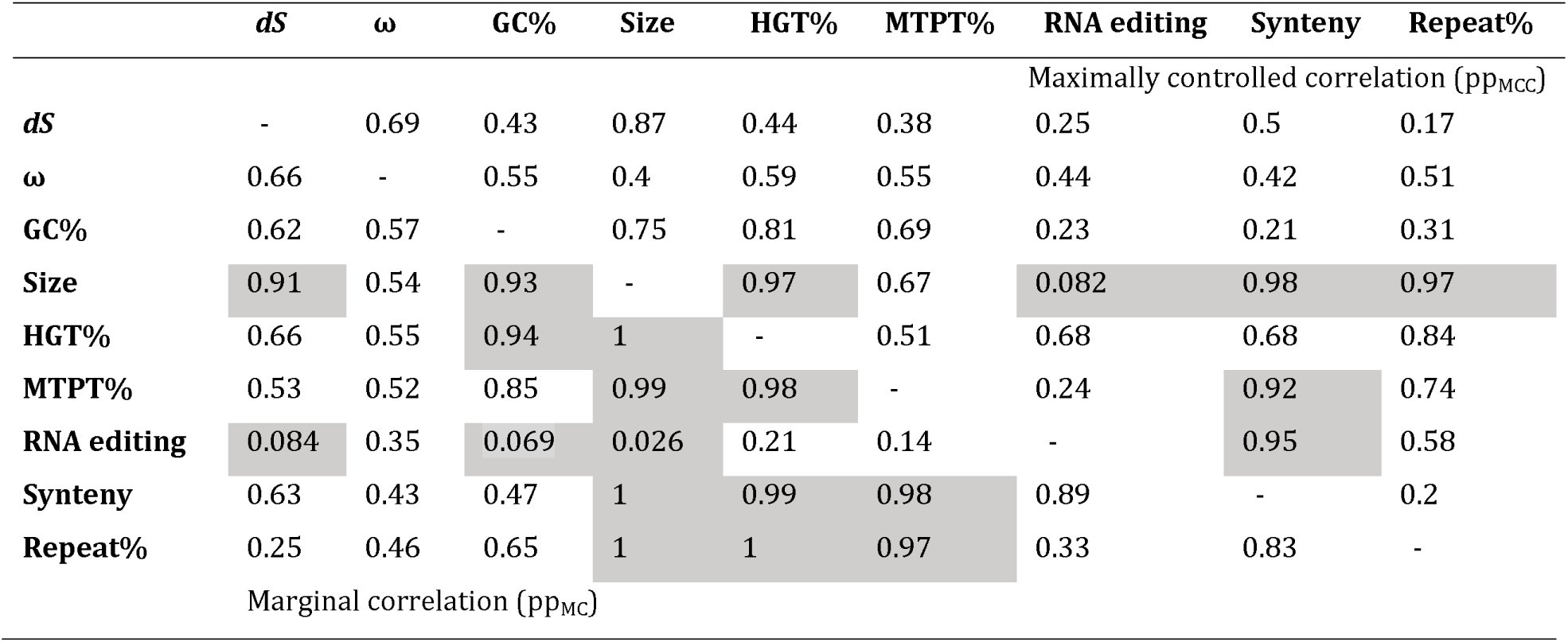
Correlation of mitochondrial genetic traits measured under the multivariate phylogenetic comparative framework implemented in COEVOL. Posterior probabilities of the marginal correlation (pp_MC_) shown below the diagonal include indirect and simultaneous correlation of the two variables, while posterior probabilities of maximally controlled correlation (pp_MCC_) shown above the diagonal control for partial correlation. Posterior probabilities (pp) toward 1 indicate a positive correlation; pp toward 0 indicates a negative correlation. Comparisons with pp values >0.9 and <0.1 are highlighted in gray.

## DISCUSSION

Orobanchaceae presents a compelling opportunity to understand the connection between mitogenome evolution and various modes of parasitic lifestyle. Unlike previous studies that overlooked noncoding regions and claimed limited influence of parasitism on mitogenome evolution, we not only revealed distinct and predictable changes in genome shuffling, RNA editing, and IGT and HGT content during the transition to holoparasitism, but also demonstrated the underlying molecular mechanisms. Overall, relaxed selection does not play the dominant role in these processes, but a rather stronger link could be made with biases in DNA recombination and repair processes, especially those involving RNA-mediated retroprocessing.

### Higher mutation rate and retroprocessing activities in holoparasites promote loss of RNA editing sites

RNA editing is a mandatory post-transcriptional correction step to restore conserved amino acids during mitochondrial gene expression across plants [41,42]. Although it has been hypothesized to buffer mutation, optimize GC content, and maintain functional and genetic variation [43–45], there is a growing consensus that RNA editing emerged through nonadaptive processes in angiosperms [46–48]. Our results strongly corroborate this nonadaptive view because (i) Orobanchaceae diversified with increasingly stringent selection against RNA editing and the free-living *Rehmannia* contains the most edited sites (n = 412; Table S7); (ii) high mutation rate is significantly linked with rapid loss of RNA editing sites, supporting its high mutational burden (PGLS *p*-value = 8.6e-4; Fig. S11)[49]; and (iii) loss of editing by C-to-T mutation is preferentially fixed and no reverse T-to-C mutations are identifiable in species impacted by retroprocessing over millions of years (i.e., the common ancestor of *Neobartsia* and *Lathraea*).

Holoparasites generally experienced more point mutations and retroprocessing that synergistically promoted loss of RNA editing sites compared to hemiparasites (Fig. 2). This might be attributed to selection — the cost of RNA editing, such as energy demands and vulnerability to mutations disrupting editing site recognition [49,50], may differ among lifestyles. Alternatively, this might be an indirect outcome of the difference in the baseline mutation and recombination rate, which we argue to be a more probable cause. First, holoparasitic species have slightly higher substitution rates in their mitochondrial genes (Fig. 4A) and hence more opportunity for spontaneous C-to-T mutations to be preferentially fixed at the editing sites (Fig. S11). Such a positive correlation between evolution rate and loss of RNA editing sites is also reported in the plant genus *Silene* and the similar rates of C-to-T substitution at synonymous versus nonsynonymous editing sites support a neutral role of these substitutions [47,51]. Second, holoparasites are more prone to retroprocessing potentially due to the additional flux of host-derived RNA and the need for retroprocessing-based DNA repair. We demonstrated that host-derived plastid RNA boosted retroprocessing of MTPT more than tenfold in holoparasites (77,602 bp vs 6,368 bp in Fig. 3B). It is reasonable that host-derived mitochondrial RNAs can similarly contribute to the loss of RNA editing sites via retroprocessing. The three independent HGTs of host *mttB* into Orobanchaceae, Cynomoriaceae, and Balanophoraceae parasites are most likely from retroprocessed host mitochondrial RNA and indirectly support this (Fig. S2). In addition, the reduced synteny and abundant repeats in holoparasites imply more active genome shuffling, greater risk of DNA damage, and hence higher demands for DNA repair. Studies have shown that reverse transcribed RNA can repair DNA double strand breaks [52], and the DNA repair process itself can promote C-to-T point mutations as well [53]. In summary, higher mutation rates and genomic shuffling likely promoted the loss of RNA editing in holoparasitic Orobanchaceae, the result of which may have indirectly resulted in streamlining mitochondrial gene expression in these parasites.

### A dosage effect mechanism for MTPT content in green plants

Intracellular gene transfers from the plastid (i.e., MTPT) comprise between 1 to 10% of the mitogenome in seed plants [54,55]. Our estimation in Orobanchaceae falls within this range (0.53–17.3%) but holoparasites exhibit drastic decline compared to hemiparasites. This general trend extends to all heterotrophic angiosperms examined in our study (Fig. 3A) and probably to parasitic algae as well due to their diminished photosynthetic capacity and degenerated plastomes. In-depth investigation revealed that such disparity is primarily driven by the sharp decrease of native MTPT in holoparasitic species, although a surge in alien MTPT slightly compensated the differences (Fig. 3B). The biased distribution of MTPT donor regions on the plastome also enabled the discovery of a novel RNA-mediated mechanism for horizontal and intracellular gene transfer.

Elucidating the molecular mechanism for highly fortuitous events like gene transfer is exceptionally challenging in eukaryotes because they are too rare to be studied *in vivo.* However, 73.7% of the alien MTPT sequences strictly reside within plastid coding regions, which is in direct contrast to native MTPTs (5.7%) and thus strongly supports their identity as reverse transcribed mRNA. This RNA-mediated mechanism was previously proposed for mitochondrion-to-nucleus gene transfers [35,56,57], but never demonstrated in cross-species and inter-organelle transfers as we have done here. Here we further argue that retroprocessing is responsible for a significant proportion of native MTPTs, and likely involves precursor RNA operons instead of mature mRNA. This stems from (i) a 7.3 kb hotspot of native MTPT in holoparasites that coincides with the *rrn* operon in the inverted repeat region and (ii) two cold zones of MTPT in hemiparasites that overlap with the *ndhC–J* and *ndhH–D* operons (Fig. 3B). These operons are bacterial-like polycistronic RNA containing multiple premature RNAs. The *rrn* operon is the most abundant transcript in plastids [58,59] whereas the *ndhC–J* and *ndhH–D* operons are amongst the rarest, showing low expression and RNA polymerase binding affinity in *Arabidopsis thaliana* [60–62]. The same hotspots and cold zones of MTPT have been reported in numerous free-living angiosperms [63,64], suggesting the universal applicability of this dosage effect mechanism to explain MTPT abundance across green plants.

On the other hand, DNA-mediated MTPT likely still dominates in most species and the processes behind its drastic decline in holoparasites are more complex. One conceivable mechanism to explain this is the “limited transfer window” hypothesis—species with a single plastid per cell experience less gene transfer because lysis of this plastid is lethal to cell [65,66]. However, in Orobanchaceae, the abundance of plastomes did not show a significant difference between hemi- and holo-parasites (phylANOVA *p-*value = 0.786) and there was no correlation between plastome abundance and MTPT content either (PGLS *p-*value = 0.635; also see caveats in METHODS). Moreover, MTPT in holoparasites declines at a rate that greatly exceeds their plastome size reduction (PGLS adjusted R^2^ = -0.04753; *p-*value = 0.45; Fig. S12), thus plastid DNA quantity alone cannot sufficiently explain the disparity.

This prompted our speculation of additional mechanisms altering plastid DNA accessibility for transfer, which we term the “resting DNA” hypothesis. This hypothesis postulates that the evolution of holoparasitism fundamentally shifts the molecular processes generating free plastid DNAs as the donor for MTPT. Replication, recombination, and lysis of the plastid all have the potential to introduce double strand breaks and free DNA fragments. While young photosynthetic leaves carry thousands of constantly replicating plastids per cell, the reproductive and meristematic cells in holoparasites contain fewer plastids that might be less active in replication [67]. These resting plastid DNAs are inaccessible for gene transfers and the biased distribution of replication origins or recombination hotspots can contribute to the uneven transferability across the genome (Fig. 3B). Along these lines, the retention of photosynthetic tissue in cryptic hemiparasites like *Harveya* and *Lathraea* facilitates the maintenance of high MTPT. Future studies can test this resting DNA hypothesis by quantifying DNA synthesis activity *in vivo* [68,69] or examining the expression of replication-related genes in species with different lifestyles.

In summary, MTPT content in plant mitochondria is maintained by both RNA- and DNA-mediated processes. The RNA-mediated pathway may recruit native precursor RNA operons or alien mature RNAs. Transition to a heterotrophic lifestyle may greatly reduce MTPT content because the genetic donors for both the DNA-mediated pathway (free plastid DNA) and the RNA-mediated pathway (precursor plastid RNA) are reduced in heterotrophs, although horizontally acquired RNA from the host may slightly compensate this phenomenon in parasitic plants.

### The limited role of selection

Unlike recombination and retroprocessing events that profoundly shaped the mitogenome landscape in holoparasites (e.g., loss of RNA editing, MTPT, and HGT), we found limited evidence for universal lifestyle-linked relaxed selection. For mitochondrial genes, both hemi- and holoparasites have similar ω values (Fig. 4B). These mitochondrial ω ranged between 0.3 to 0.5, which were even higher than those from the degraded plastid genome in holoparasites (ω = 0.2­–0.3; Fig. 4B). This result could indicate overall relaxation in plant mitochondria regardless of lifestyle, despite the conventional view of their conserved sequence and function. Genes including *ccm, cob,* and *mttB* showed significant relaxed selection in holoparasitic lineages, but the clades involved in selective shifts varied depending on the gene (Table 2). The loss of mitochondrial-encoded *mttB* in the holoparasitic *Mannagettaea* may also stem from relaxed selection (Fig. S2). Loss of *mttB* is uncommon among angiosperms and is only documented in the parasitic mistletoe *Viscum* and the mycoheterotrophic *Epirixanthes* [12,27], which may suggest convergent relaxed selection on its retention in the mitogenome among heterotrophic plants. On the other hand, intensified selection was found in NADPH dehydrogenase and cytochrome c oxidase among holoparasites. Therefore, the occurrence and direction of selective shifts on mitochondrial genes is highly gene- and clade-specific, which may explain the lack of overall signals in previous investigations when all mitochondrial genes were concatenated and examined [10].

In addition to coding regions, integration of excessive non-coding DNA such as MTPTs and HGTs increases the mutational burden of the mitogenome [49], which may be interpreted as a result of relaxed selection. For the plastid genome, relaxed selection following parasitism has long been recognized to drive the concerted acceleration of evolutionary rates and genomic rearrangements in plastomes [5]. In mitochondria, nucleotide substitution parameters including *d_S_* and GC content were correlated with many genomic features (Table 2), but selective constraints (ω) did not show correlation with any substitutional or genomic traits (pp_MC_ = 0.32–0.68). Moreover, mitogenome size was positively correlated with *d_S_* (Table 2), which is consistent with the trend reported in *Silene* [70]. However, such a positive correlation is at odds with the prediction from the mutation burden hypothesis where high mutation rates create selection against large genomes [49].

Instead, mitogenome size scales strongly with traits directly or indirectly associated with recombination and repair processes, including MTPT, HGT, repeats, GC content, and RNA editing (pp_MC_ =0.91–1; pp_MCC_ = 0.81–0.98). In particular, GC content and RNA editing are not directly correlated with genome size in PGLS regressions (*p-*value = 0.883), but only show negative correlation under the multivariate comparative framework (pp_MC_ = 0.069; pp_MCC_ = 0.082). This might result from their indirect correlation to a third, unmeasured trait—recombination. Homologous recombination is known to promote GC-biased gene conversion, loss of RNA editing via retroprecessing, and integration of alien DNA that bloats the genome [71,72]. In *Silene* species with exceptionally large mitogenomes, high recombination rates are also responsible for rapidly evolving genome structure with numerous extra chromosomes that eventually inflate the mitogenome [70,73,74]. Given the rapid turnover rate of mitogenomes in holoparasites especially (e.g., nearly complete lack of synteny and MTPT purging), recombination emerges as a more profound force shaping genome evolution compared to selection.

### A roadmap for mitochondrial genome evolution in heterotrophic plants

To summarize our results and provide an integrative framework for future studies, we propose a universal roadmap of plant mitogenome evolution in response to parasitism (Fig. 5). This model can be applied to other heterotrophic lineages including algae and mycoheterotrophic plants. Many macrostructural changes in the mitogenome, such as MTPT and HGT, follow a predictable path after the establishment of parasitism; selection and substitution processes, on the other hand, exhibit lineage- and gene-specific patterns.

**Figure 5.**
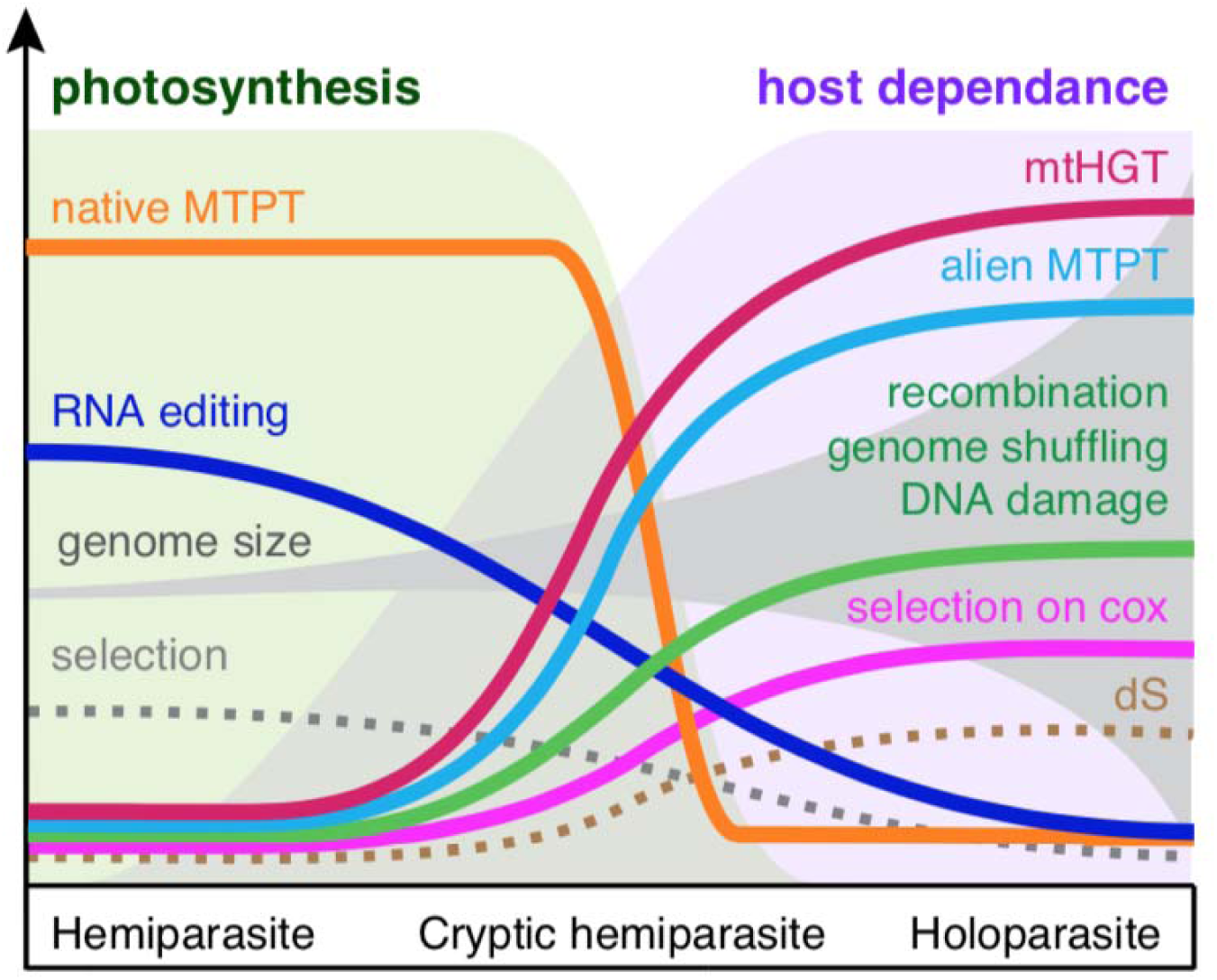
Roadmap of mitogenome evolution in parasitic plants. The model illustrates the evolution of genomic traits and evolutionary rates during the transition from hemiparasites with high photosynthetic capacity to cryptic hemiparasites with rapidly declining photosynthesis, and holoparasites showing complete host dependence. Most genomic traits follow a predictable path. This includes the rapid decline of native mitochondrial plastid DNAs (native MTPT) after the loss of photosynthesis, gradual loss of RNA editing, increased uptake of mitochondrial horizontal gene transfer (mtHGT) and alien MTPT, all of which are directly or indirectly associated with genomic recombination, shuffling, and DNA damage repair processes. Changes in evolutionary rates (*d_S_*) and selection are less predictable and display lineage and gene specific patterns. We predict a subtle relaxation in the overall mitochondrial function and slight increase in *d_S_*. But the cytochrome c oxidase (*cox*) genes are likely to experience intensified selection in holoparasites with high metabolic rates. Genome size similarly contains substantial contingence where alien genetic material like mtHGT may inflate the genome size, but increased recombination can rapidly remove long stretches of mitogenome as well.

Parasitism establishes direct vascular connections with other plant species, which increases the load of alien RNA and DNA molecules that facilitate gene transfer to the mitogenome. These transfers come from the plastid, mitochondrial, and even nuclear genomes of hosts and are of minimal influence in hemiparasites. Besides host-derived genetic materials, native sequences from the plastome frequently enter the mitogenome via RNA- and DNA-mediated pathways. In free-living and hemiparasitic species, integration of native MTPT often involves reverse transcription of precursor RNA operons and is strongly linked with transcript abundance. The loss of photosynthesis in holoparasites marks a major transition in the dynamics of gene transfers. The greatly diminished photosynthetic capacity decimates the transcription of most plastid genes as well as their entrance to the mitogenome via RNA-mediated retroprocessing. The only plastome region being actively transferred in holoparasites involves the rRNAs, whose function seems indispensable in all plants with a plastome [75–77]. The enhanced hydraulic and nutritional reliance on hosts, on the other hand, significantly increases the flux of host-derived DNA and RNA, leading to accelerated rates of HGT. Cryptic hemiparasites with photosynthetic capacity and high host dependence will display both high MTPT and HGT content. Among various mechanisms of gene transfers, retroprocessing of mature mRNA from the host plastome increases alien MTPT in holoparasites and similar processes involving mitochondrial genes result in the loss of RNA editing and introns. This error-prone retroprocessing may also contribute to the increased gene substitution rate often seen in holoparasites [78]. These insertions introduce homologous regions in the mitogenome that may eventually become repeats with recombination capacity (e.g., 9 parallel origins of structural repeats involving nad-like genes; Table S6). In addition, loss or alteration of nuclear-encoded mitochondrial DNA replication, recombination, and repair (RRR) machinery may also influence organellar nucleotide substitution. We recently discovered that species in the holoparasitic tribe Orobancheae have lost 20% mitochondrial RRR gene compared to other angiosperms [19]. Specifically, the plant mitochondrial recombination surveillance protein *RecA3* was lost in four out of five Orobancheae species examined and its disruption resulted in extensive rearrangement of the mitochondrial genome in *Arabidopsis* [79]. All of these factors may synergistically contribute to more frequent genome shuffling and DNA damage in holoparasites.

These excessive genome structural changes are deleterious because they can disrupt gene expression (e.g., pseudogenization of *mttB* due to HGT; Fig. S2). However, the overall ineffective or relaxed selection in the mitochondria permits the proliferation of these deleterious mutations. Some genes including *ccm, cob,* and *mttB* show significant relaxed selection in holoparasitic lineages. The precise causes of relaxed selection in mitochondrial genes remain to be explored but may be linked to the reliance on external carbon or altered energy demands of stomata and hydraulic regulation [80]. Conversely, intensified selection is found in cytochrome c oxidase among holoparasites. Adaptive changes in cytochrome c oxidase can boost mitochondrial respiratory efficiency for species with high metabolic rates, which is demonstrated in the carnivorous bladderwort *Utricularia* [81] and potentially relevant for holoparasites like *Lathraea* as well because they actively pump water to sequester nutrients from hosts [38].

Despite these general trends, mitogenome evolution in heterotrophic species is subject to substantial contingency. This originates from their altered energy metabolism that may either increase (e.g., thermal genesis in *Rafflesia*) or decrease (e.g., glycolysis-based respiration in mistletoe) metabolic demands depending on the lineage [82,83]. In regard to the mitogenome structure, parasitism introduces alien sequences that bloat the genome size, but also increases recombination that can rapidly remove non-essential DNA [84]. Holoparasitic species with high metabolic rates are prone to reactive oxygen species stress and the deletion-biased DNA repair mechanism would lead to genome downsizing as well [85,86]. The complex interplay of these molecular processes thus generates the fascinating diversity of genome structure in heterotrophic plants ranging from the megabase-sized multi-chromosome genome of *Cistanche* to the miniature mitogenome of *Rhopalocnemis* (Balanophoraceae).

## METHODS

### Taxon sampling and DNA sequencing

Our taxon sampling included 45 representative species from 28 (28%) genera in Orobanchaceae (Table S1). Specifically, we included all three independent origins of holoparasitism in the tribe Orobancheae, the Hyobanche clade, and *Lathraea*. We generated genome sequences for 32 species and obtained public data for the other 13 species from NCBI GenBank as reads or assemblies. DNAs from all but two species were extracted from herbarium specimens, dating back as early as 1922 (Table S1). To prepare for DNA extraction, plant materials were flash-frozen in liquid nitrogen in 2 ml tubes and immediately homogenized using the 2010-115 Geno/Grinder High-Throughput Homogenizer (SPEX SamplePrep LLC, NJ, USA). Subsequent DNA extraction followed the standard CTAB protocol [87,88]. Precipitated DNAs were dissolved in TE buffer and then cleaned using AMPure XP magnetic beads (Agencourt, MA, USA) following the manufacturer’s instructions. Final DNA quality and concentration was assessed using Nanodrop 2000/2000c Spectrophotometer and Qubit Fluorometric Quantification (Thermo Fisher Scientific, MA, US). Due to the presence of orobanchoside and other chemicals generated by specimen preparation [89], the success rate of DNA extraction from herbarium specimens was especially low for hemiparasites (∼20%) compared to holoparasites (∼80%). Finally, DNA samples passing basic quality controls were shipped on dry ice to BGI USA (San Jose, CA, USA) for library preparation and sequencing on the DNBSeq platform (MGI Tech, Shenzhen, China). An average of 8.3 giga base pairs (bp) of 150-bp paired-end reads were generated for each species. All newly generated read data were deposited at GenBank under BioProject PRJNA1169115.

### Genome assembly and annotation

We trimmed adapters and filtered low-quality reads using TrimGalore v.0.5.0 under the default settings [90] on the high performance computing cluster hosted at the Texas Advanced Computing Center. Organellar genome assemblies were conducted using GetOrganelle v1.7.7.0 [91]. First, we used the built-in database and the default kmer sizes to assemble the plastid genomes for all species. We generated complete plastomes for all species with a median base coverage of 1,287× (Table S3). These plastome assemblies were subsequently manually inspected and circularized in Bandage [92]. Second, we removed plastid reads for mitogenome assembly by mapping reads to the corresponding plastid assembly using the end-to-end algorithm implemented in bowtie v2.5.1 [93]. The resulting sam files were filtered through a custom Python script to remove reads with 100% sequence similarity and >98% sequence overlap with the plastome assemblies (all scripts available on GitHub https://github.com/lmcai/Orobanchaceae_comparative_mitome). This threshold allowed us to retain potential mitochondrial plastid DNA (MTPT) in a conservative manner, and exclude any false MTPTs with very high sequence identity to the plastome. The filtered reads were used for mitogenome assembly in GetOrganelle with the recommended parameter settings (-R 20 -k 65,85,105 -P 1000000). The resulting assembly graph was visualized in Bandage and manually circularized when possible. Reads were mapped back to the final assemblies using BWA-MEM 0.7.17 [94] and visualized in Tablet to correct assembly errors [95].

As a result, we completed mitogenomes of 24 species with fully connected De Bruijn assembly graphs in networks (Table S4). Eleven species have various unconnected edges in their assemblies, likely caused by missing data or linearly branched genome structure [23]. Given their high base coverage (> 128×) and complete mitochondrial gene sets, we consider these assemblies as nearly complete.

Annotation of organellar genomes was conducted using the annotation transfer function in Geneious Prime 2019.1.3 and manually inspected (Biomatters Ltd, Auckland, New Zealand, http://www.geneious.com/). The plastid (GenBank ID: NC_034308) and mitochondrial (GenBank ID: OM397952) annotations from *Rehmannia glutinosa* were used as references. Annotated exons were mapped back to the assembly to identify trans-spliced introns. These careful inspections of assembly and annotation revealed several assembly errors in published mitogenomes, which we corrected later by read mapping (e.g., incorrect frameshifts in *atp1, cox3*, and *nad5* in *Cistanche salsa* GenBank ID ON890402–ON890407; see notes in Table S4).

### Relative copy numbers of organellar genomes

To explore the abundance of organellar genomes within a plant cell, we characterize the copy number of mitogenomes, plastomes, and nuclear genomes based on their read base coverage. One important caveat is that there is great variation in the number of genomes carried by each organelle ranging from dozens to hundreds depending on the species and cell type [96]. Thus these copy numbers may not reflect the abundance of organelles. Here, we only used our sequencing data because the libraries were built with an unbiased sampling process. To calculate base coverage for organelle genomes, raw reads were mapped to the assemblies using BWA-MEM. The base coverage for each site was calculated from the resulting bam file using the ‘samtools depth’ function in samtools. We also used bowtie2 for read mapping to confirm that the results were similar and were thus robust to mapping algorithms and parameters (Table S3). The coverage reported in the Results was inferred from BWA-MEM only. Finally, the nuclear genome coverage was estimated by kmer distribution using Jellyfish v2.3.0 with a kmer size of 21 [97].

### Phylogeny reconstruction and time tree inference

We reconstructed mitochondrial and plastid phylogenies with genes and conserved introns. For the mitochondrial phylogeny, 31 protein-coding genes, 3 rRNAs, and their conserved introns (Table S11) were aligned using MAFFT-linsi and manually inspected in Geneious. We then inferred a Maximum Likelihood phylogeny based on the concatenated loci in IQ-TREE v2.2.2.6 [98]. A gene-by-gene partition was applied and the best substitution model was determined by IQ-TREE. Branch support was evaluated by 1000 ultrafast bootstrap replications (UFBP). The plastid-based phylogeny was inferred using a similar pipeline with a subset of 25 plastid genes with conserved sequences in holoparasites [5] (Table S12).

An ultrametric time tree was inferred for downstream comparative analyses (Fig. S13). To accomplish this, we used the divergence times estimated by Mortimer et al. (2022) to fix the ages of six major nodes in the mitochondrial phylogeny (Fig. S13). We then used the penalized likelihood implemented in TreePL v1.0 [99] to generate the time tree. TreePL was run three times to prime and cross validate the analyses. Divergence time was subsequently estimated with an optimum smoothing parameter of 1000.

### Repeat annotation

We used two methods to localize and quantify repeats in mitochondrial genomes. First, we used the BLAST-based tool ROUSFinder v2.0 [100] to identify interspersed repeats longer than 17 bp (-m 17). Then the total size of the annotated repeats was calculated by bedtools merge v2.18 [101].

Secondly, we used a custom Python script (de_bruijn_graph_based_repeat_identification.py, see GitHub for details) to identify repeats bridging multiple assembly contigs. Here, repeats are identified as contigs with four or more connections with neighboring contigs and a higher kmer coverage that aligns with the number of connections (e.g., 4 connections = 2× coverage; Fig. S14). This graph-based repeat identification is more robust because it is supported by reads flanking adjacent contigs. To characterize the homology of these repeats, we conducted all-by-all BLAST of the repeat sequences and then clustered them into network modules using the cluster_louvain function in the R package igraph, which implements the multi-level modularity optimization algorithm to define community structure [102].

### Genome synteny assessment

Pairwise genome alignment was conducted in Mummer v4.0.0.0 [103] using the mitogenome of *Rehmannia glutinosa* as the reference. To accommodate the large variation of mitogenomes, Mummer was applied with 1000 bp maximum gap length (-g 1000), 1000 bp maximum alignment extension (-b 1000), and using all maximal exact matches (--maxmatch). The resulting alignments were visualized using the mummerCoordsDotPlotly.R function in dotPlotly (https://github.com/tpoorten/dotPlotly). The coordinates of the aligned regions were then extracted and summarized using the ‘bedtools coverage’ to identify genomic regions that were universally conserved (see GitHub).

### RNA editing

We used a neural network based software Deepred-Mt [104] to predict RNA editing in Orobanchaceae. This tool is trained from an extensive set of experimentally verified RNA editing sequences in plant mitochondria and is demonstrated to be more accurate than similarity-based tools [104]. Here, we set a minimum threshold of 0.9 probability score for RNA editing, which resulted in <0.1% type I error in the training dataset (Fig. S15). We then used a custom script mapping_deepredmt_to_alignment.py (available on GitHub) to flag predicted RNA editing sites and realign these sequences to investigate their evolutionary trajectory. Ancestral states of these RNA editing sites were determined under the mitochondrial phylogeny based on the Maximum Parsimony criteria implemented in the MPR function from the ape R package [105,106].

### MTPT annotation

To accurately identify MTPTs, we generated a comprehensive plastome database that included both highly divergent sequences from holoparasites and conserved sequences from representative free-living angiosperms from the host lineage (Table S13). This allowed us to identify MTPT sequences from degraded holoparasitic plastids as well as MTPTs horizontally transferred from their hosts.

To calculate the proportion of MTPT, each mitogenome was searched against the plastome database with a stringent e-value threshold of 1e-70 in BLAST. All hits were consolidated using ‘bedtools merge’ to summarize their total length. Five additional non-photosynthetic angiosperm lineages were sampled for comparative purposes, including Balanophoraceae (Santalales), Hydnoraceae (Pipeales), Cynomoriaceae (Saxifragales), *Cuscuta* (Solanales), and the mycoheterotrophic *Monotropa* (Ericales) (Table S8). Within each lineage, mitogenomes from multiple photosynthetic and non-photosynthetic members were examined for their MTPT content. A mitochondrial phylogeny was generated for these species using the same set of 24 core genes in Table S7 in IQ-TREE. An ultrametric tree was then inferred using the penalized likelihood method implemented in the chonos function in the R package ape [106](Fig. S16). The phylogenetic ANOVA test implemented in phytools was then used to test the correlation between MTPT abundance and life history strategy.

To further characterize the evolutionary history of MTPT, we reconstructed the phylogeny for all 1,094 MTPT fragments identified across Orobanchaceae species. To do this, a custom Python program HGTscanner_mtpt.py (available on GitHub) was generated to establish sequence homology, infer phylogeny, and identify intra- or inter-species gene donors. Briefly, the NCBI nucleotide (NT) database was queried (accessed 10 November 2023) for each MTPT to identify homologous plastid regions. A phylogeny was subsequently inferred using IQ-TREE v2.2.2.7 [98] and the sister lineage was reported. Detailed program and parameter descriptions can be found in Supplemental Note 2.

### HGT annotation

To accurately identify mitochondrial HGTs, we developed a Python tool HGTscanner to precisely characterize the location, donor, and recipient of alien genetic fragments (Fig. S17). Here, the two main challenges lie in the difficulty of (1) establishing homology across highly dynamic mitogenomes, especially non-coding regions; and (2) identification of HGT donors from dozens of potential host families while taking into account phylogenetic uncertainty.

To mitigate these challenges, HGTscanner first masked exons and MTPT in the query mitogenome assembly and then BLASTed it against the entire Viridiplantae mitochondrial sequence database from NCBI GenBank (70,706 records; accessed 10 November 2023). We found such masking to be essential for avoiding the large number of BLAST hits in conserved coding regions and MTPTs. These BLAST hits were then ordered based on location and consolidated into longer synteny blocks for downstream phylogenetic analyses. Each synteny block was further divided if more than 50% of the BLAST hits consisted of multiple genomic regions (Fig. S17). Such finer division is essential for the identification of shorter HGTs nested within long synteny blocks (e.g., alien intron nested within native exons in *cox2*). Sequences from each synteny block were subsequently aligned using the MAFFT-einsi algorithm [107]. A maximum likelihood phylogeny was inferred for each block by IQ-TREE with 1000 ultrafast bootstrap replicates.

HGT was evaluated based on stringent criteria using BLAST and phylogenetic evidence. Briefly, a genetic locus was classified as “high-confidence HGT” if (1) Sequences with high identity (BLAST e-value < 1e-20) are found in Orobanchaceae and one non-Lamiales family (BLAST-based evidence); (2) Sequences with high identity (BLAST e-value < 1e-20) are found in only one Orobanchaceae species and other land plant families (BLAST-based evidence); (3) the target Orobanchaceae parasite is nested well within a non-Lamiales family with >85 UFBP support (phylogeny-based evidence). The other scenarios are variously classified as ‘VGT’ (vertical gene transfer), ‘putative HGT’, or ‘inconclusive’ (Fig. S17). These stringent criteria are effective in removing incorrect identification of HGT due to phylogenetic uncertainty, but may suffer from high false negative rates because most HGTs within Lamiales, especially from viable hosts in Lamiaceae and Plantaginaceae, will be classified as VGT. Detailed program parameters and software pipeline are described in Supplementary Note 2. Downstream comparative analyses on HGT were based on high-confidence HGT only.

### Molecular evolutionary rates

We used various models to investigate mitochondrial selection at the species and gene levels. To prepare the input sequences, coding sequences from mitochondrial genes were aligned with the codon-aware aligner MASCE v2.01b [108]. The resulting alignments were verified against the TAIR database (www.arabidopsis.org) to ensure the correct reading frames. Sequences containing premature stop codons and frameshifts were removed. All RNA-editing sites were masked prior to subsequent analyses.

The overall non-synonymous (*d_N_*) and synonymous substitution rates (*d_S_*) were calculated by pairwise comparison to the free-living outgroup *Rehmannia glutinosa* using concatenated plastid or mitochondrial coding sequences in CODEML v4.10.7 [109]. To more finely characterize selection on individual branches, we also inferred the substitution rates under the free-ratio model in CODEML (model = 2). For the RELAX analysis implemented in HYPHY 2.5.33 [110], we grouped genes into functional groups (Table S10) and then tested for relaxed or intensified purifying selection for each life history group. The RELAX analyses were conducted under the GTR substitution model and assuming three rate classes across sites. The mitochondrial phylogeny was used as the guidance tree and we tested three scenarios of selective shifts in foreground branches: (i) holoparasitic tribe Orobancheae; (ii) all three holoparasitic lineages (and their internal branches); and (iii) all parasitic Orobanchaceae. The best model was determined by the corrected Akaike Information Criterion (AICc). The branch model (i) that tested for differentiated selection in tribe Orobancheae was based on our investigation of nuclear-encoded mitochondrial genes where tribe Orobancheae showed unique and exceptional gene losses not seen in the other two holoparasitic lineages [19]. The selection parameter *k* inferred from the general descriptive model was used for visualization in Fig. 4 using a custom python script ‘hyphy_json_parsser.py’ (available on GitHub).

### Phylogenetic comparative hypothesis testing

Most tests of lifestyle related shifts of genetic traits were conducted using phylogenetic ANOVA (phylANOVA) or phylogenetic generalized least squares (PGLS). The ultrametric species tree inferred from the concatenated Orobanchaceae mitochondrial genes was used as the reference phylogeny. Species were classified based on their photosynthetic capacity as “holoparasites” or “photosynthetic” (including cryptic hemiparasites, hemiparasites and free-living species). For phylANOVA analyses, the PhylANOVA function from the R package phytools was used to perform hypothesis testing using 1000 simulations, posthoc tests to compare the mean among groups, and the “holm” method to adjust *p*-values to account for multiple testing [111]. For PGLS analyses, the comparative.data function from the R package caper [112] was used to prepare the comparative dataset with phylogeny, trait value (e.g., HGT content), and group assignment. The pgls function from caper was subsequently applied to perform the PGLS regression.

Correlation of genomic traits (size, repeats, synteny, RNA editing, MTPT, HGT) and nucleotide substitution (*d_S_*, ω, GC%) were analyzed with COEVOL v.1.6 [40]. COEVOL uses Bayesian inference and MCMC methods to fuse phylogenetic substitution models with multivariate Brownian comparative models. COEVOL was run with two chains, each sampling every ten points until 10,000 samples were collected. Convergence was verified in Tracer v1.7.2 [113] and the two chains were merged with 10% burn-in. The posterior probability and covariance matrix were summarized using the readcoevol command in COEVOL.

## ACKNOWLEDGEMENTS

We thank Thomas E Juenger for sharing the Genogrinder equipment for tissue homogenizing and DNA extraction. We thank TEX-LL, HUH, and NYBG for allowing us to perform destructive sampling of herbarium specimens. This work is supported by the Stengl Wyer Postdoctoral Fellowship from the University of Texas at Austin, Texas Ecological Laboratory Program, and the Fellowship in Plant Science Research from the Oak Spring Garden Foundation to LC; NIH grant R35GM142836 to JCH; Texas Ecological Laboratory Program and Sidney F. and Doris Blake Professorship in Systematic Botany from the University of Texas at Austin to RKJ.

## DATA AVAILABILITY

All newly generated raw sequence data were deposited at GenBank under BioProject PRJNA1169115 (https://www.ncbi.nlm.nih.gov/sra/PRJNA1169115) and the SRA accession numbers were provided in Table S1. The assemblies of newly generated mitogenomes and plastomes were deposited at GenBank ###. The code used for data processing and analysis is openly available on GitHub at https://github.com/lmcai/Orobanchaceae_comparative_mitome. The genome assemblies, annotations, gene alignments, and phylogenies supporting the findings of this study including RNA editing, molecular evolution, and gene transfers are available as supplementary data deposited in the Zenodo Digital Repository (https://doi.org/10.5281/zenodo.14862040).

## AUTHOR CONTRIBUTIONS

Conceptualization: LC, RKJ; Funding Acquisition: LC; Data Curation: LC; Formal Analysis: LC; Methodology: LC; Supervision: RKJ, JCH; Writing – Original Draft Preparation: LC; Writing – Review & Editing: LC, RKJ, JCH.

## REFERENCE

1. John U, Lu Y, Wohlrab S, Groth M, Janouškovec J, Kohli GS, et al. An aerobic eukaryotic parasite with functional mitochondria that likely lacks a mitochondrial genome. Sci Adv. 2019;5: eaav1110. doi:10.1126/sciadv.aav1110

2. Yahalomi D, Atkinson SD, Neuhof M, Chang ES, Philippe H, Cartwright P, et al. A cnidarian parasite of salmon (Myxozoa: Henneguya) lacks a mitochondrial genome. Proc Natl Acad Sci. 2020;117: 5358–5363. doi:10.1073/pnas.1909907117

3. Karnkowska A, Vacek V, Zubáčová Z, Treitli SC, Petrželková R, Eme L, et al. A eukaryote without a mitochondrial organelle. Curr Biol. 2016;26: 1274–1284. doi:10.1016/j.cub.2016.03.053

4. Těšitel J. Functional biology of parasitic plants: a review. Plant Ecol Evol. 2016;149: 5–20. doi:10.5091/plecevo.2016.1097

5. Wicke S, Müller KF, Quandt D, Bellot S, Schneeweiss GM. Mechanistic model of evolutionary rate variation en route to a nonphotosynthetic lifestyle in plants. Proc Natl Acad Sci. 2016; 201607576.

6. Wicke S, Müller KF, de Pamphilis CW, Quandt D, Wickett NJ, Zhang Y, et al. Mechanisms of functional and physical genome reduction in photosynthetic and nonphotosynthetic parasitic plants of the broomrape family. Plant Cell. 2013;25: 3711–3725.

7. Wolfe AD, dePamphilis CW. The effect of relaxed functional constraints on the photosynthetic gene rbcL in photosynthetic and nonphotosynthetic parasitic plants. Mol Biol Evol. 1998;15: 1243–1258. doi:10.1093/oxfordjournals.molbev.a025853

8. Sanchez-Puerta MV, Ceriotti LF, Gatica-Soria LM, Roulet ME, Garcia LE, Sato HA. Invited Review Beyond parasitic convergence: unravelling the evolution of the organellar genomes in holoparasites. Ann Bot. 2023;132: 909–928. doi:10.1093/aob/mcad108

9. Skippington E, Barkman TJ, Rice DW, Palmer JD. Miniaturized mitogenome of the parasitic plant Viscum scurruloideum is extremely divergent and dynamic and has lost all nad genes. Proc Natl Acad Sci. 2015;112: E3515–E3524.

10. Zervas A, Petersen G, Seberg O. Mitochondrial genome evolution in parasitic plants. BMC Evol Biol. 2019;19: 87.

11. Fan W, Zhu A, Kozaczek M, Shah N, Pabón-Mora N, González F, et al. Limited mitogenomic degradation in response to a parasitic lifestyle in Orobanchaceae. Sci Rep. 2016;6: 36285.

12. Petersen G, Darby H, Lam VKY, Pedersen HÆ, Merckx VSFT, Zervas A, et al. Mycoheterotrophic Epirixanthes (Polygalaceae) has a typical angiosperm mitogenome but unorthodox plastid genomes. Ann Bot. 2019;124: 791–807. doi:10.1093/aob/mcz114

13. Haraguchi Y, Sasaki A. Host-parasite arms race in mutation modifications: indefinite escalation despite a heavy load? J Theor Biol. 1996;183: 121–137. doi:10.1006/jtbi.1996.9999

14. Bromham L, Cowman PF, Lanfear R. Parasitic plants have increased rates of molecular evolution across all three genomes. BMC Evol Biol. 2013;13: 1–11.

15. Havird JC, Trapp P, Miller CM, Bazos I, Sloan DB. Causes and consequences of rapidly evolving mtDNA in a plant lineage. Genome Biol Evol. 2017;9: 323–336. doi:10.1093/gbe/evx010

16. Healy TM, Burton RS. Strong selective effects of mitochondrial DNA on the nuclear genome. Proc Natl Acad Sci. 2020;117: 6616–6621. doi:10.1073/pnas.1910141117

17. Medeiros DB, Aarabi F, Martinez Rivas FJ, Fernie AR. The knowns and unknowns of intracellular partitioning of carbon and nitrogen, with focus on the organic acid-mediated interplay between mitochondrion and chloroplast. J Plant Physiol. 2021;266: 153521. doi:10.1016/j.jplph.2021.153521

18. Cai L. Rethinking convergence in plant parasitism through the lens of molecular and population genetic processes. Am J Bot. 2023;110: e16174. doi:10.1002/ajb2.16174

19. Cai L, Jansen RK, Havird JC. Altered mitochondrial respiration is associated with loss of nuclear-encoded OXPHOS genes in parasitic broomrapes. bioRxiv; 2025. p. 2025.02.10.637464. doi:10.1101/2025.02.10.637464

20. Ghifari AS, Saha S, Murcha MW. The biogenesis and regulation of the plant oxidative phosphorylation system. Plant Physiol. 2023;192: 728–747. doi:10.1093/plphys/kiad108

21. Havird JC, Noe GR, Link L, Torres A, Logan DC, Sloan DB, et al. Do angiosperms with highly divergent mitochondrial genomes have altered mitochondrial function? Mitochondrion. 2019;49: 1–11.

22. Weaver RJ, Carrion G, Nix R, Maeda GP, Rabinowitz S, Iverson EN, et al. High mitochondrial mutation rates in Silene are associated with nuclear-mediated changes in mitochondrial physiology. Biol Lett. 2020;16: 20200450.

23. Smith DR, Keeling PJ. Mitochondrial and plastid genome architecture: reoccurring themes, but significant differences at the extremes. Proc Natl Acad Sci. 2015;112: 10177–10184.

24. Cusimano N, Renner SS. Sequential horizontal gene transfers from different hosts in a widespread Eurasian parasitic plant, Cynomorium coccineum. Am J Bot. 2019;106: 679–689.

25. Xi Z, Wang Y, Bradley RK, Sugumaran M, Marx CJ, Rest JS, et al. Massive mitochondrial gene transfer in a parasitic flowering plant clade. PLOS Genet. 2013;9: e1003265.

26. Yu R, Sun C, Zhong Y, Liu Y, Sanchez-Puerta MV, Mower JP, et al. The minicircular and extremely heteroplasmic mitogenome of the holoparasitic plant *Rhopalocnemis phalloides*. Curr Biol. 2022;32: 470–479.e5. doi:10.1016/j.cub.2021.11.053

27. Petersen G, Cuenca A, Møller IM, Seberg O. Massive gene loss in mistletoe (Viscum, Viscaceae) mitochondria. Sci Rep. 2015;5: 1–7.

28. Lynch M. The origins of genome architecture. 1st ed. Sunderland, MA: Sinauer Associates; 2007.

29. Sanchez-Puerta MV, García LE, Wohlfeiler J, Ceriotti LF. Unparalleled replacement of native mitochondrial genes by foreign homologs in a holoparasitic plant. New Phytol. 2017;214: 376–387.

30. Shtratnikova VY, Schelkunov MI, Penin AA, Logacheva MD. Mitochondrial genome of the nonphotosynthetic mycoheterotrophic plant Hypopitys monotropa, its structure, gene expression and RNA editing. PeerJ. 2020;8. doi:10.7717/peerj.9309

31. Yu R, Chen X, Long L, Jost M, Zhao R, Liu L, et al. De novo assembly and comparative analyses of mitochondrial genomes in Piperales. Genome Biol Evol. 2023;15: evad041. doi:10.1093/gbe/evad041

32. Štorchová H, Krüger M. Methods for assembling complex mitochondrial genomes in land plants. J Exp Bot. 2024;75: 5169–5174. doi:10.1093/jxb/erae034

33. McNeal JR, Bennett JR, Wolfe AD, Mathews S. Phylogeny and origins of holoparasitism in Orobanchaceae. Am J Bot. 2013;100: 971–983.

34. Wicke S, Naumann J. Molecular evolution of plastid genomes in parasitic flowering plants. Adv Bot Res. 2018;85: 315–347.

35. Adams KL, Palmer JD. Evolution of mitochondrial gene content: gene loss and transfer to the nucleus. Mol Phylogenet Evol. 2003;29: 380–395.

36. Mortimer SM, Boyko J, Beaulieu JM, Tank DC. Synthesizing existing phylogenetic data to advance phylogenetic research in Orobanchaceae. Syst Bot. 2022;47: 533–544.

37. Randle CP, Wolfe AD. The evolution and expression of RBCL in holoparasitic sister-genera Harveya and Hyobanche (Orobanchaceae). Am J Bot. 2005;92: 1575–1585. doi:10.3732/ajb.92.9.1575

38. Atkinson MD, Atkinson E. Biological flora of the British Isles: Lathraea clandestina. J Ecol. 2020;108: 2145–2168. doi:10.1111/1365-2745.13473

39. Wertheim JO, Murrell B, Smith MD, Kosakovsky Pond SL, Scheffler K. RELAX: Detecting Relaxed Selection in a Phylogenetic Framework. Mol Biol Evol. 2015;32: 820–832. doi:10.1093/molbev/msu400

40. Lartillot N, Poujol R. A phylogenetic model for investigating correlated evolution of substitution rates and continuous phenotypic characters. Mol Biol Evol. 2011;28: 729–744. doi:10.1093/molbev/msq244

41. Gualberto JM, Lamattina L, Bonnard G, Weil JH, Grienenberger JM. RNA editing in wheat mitochondria results in the conservation of protein sequences. Nature. 1989;341: 660–662. doi:10.1038/341660a0

42. Small ID, Schallenberg-Rüdinger M, Takenaka M, Mireau H, Ostersetzer-Biran O. Plant organellar RNA editing: what 30 years of research has revealed. Plant J. 2020;101: 1040–1056. doi:10.1111/tpj.14578

43. Börner GV, Yokobori S, Mörl M, Dörner M, Pääbo S. RNA editing in metazoan mitochondria: staying fit without sex. FEBS Lett. 1997;409: 320–324. doi:10.1016/s0014-5793(97)00357-8

44. Jobson RW, Qiu Y-L. Did RNA editing in plant organellar genomes originate under natural selection or through genetic drift? Biol Direct. 2008;3: 43. doi:10.1186/1745-6150-3-43

45. Tillich M, Lehwark P, Morton BR, Maier UG. The evolution of chloroplast RNA editing. Mol Biol Evol. 2006;23: 1912–1921. doi:10.1093/molbev/msl054

46. Edera AA, Gandini CL, Sanchez-Puerta MV. Towards a comprehensive picture of C-to-U RNA editing sites in angiosperm mitochondria. Plant Mol Biol. 2018;97: 215–231. doi:10.1007/s11103-018-0734-9

47. Mower JP. Modeling Sites of RNA Editing as a Fifth Nucleotide State Reveals Progressive Loss of Edited Sites from Angiosperm Mitochondria. Mol Biol Evol. 2008;25: 52–61. doi:10.1093/molbev/msm226

48. Shields DC, Wolfe KH. Accelerated evolution of sites undergoing mRNA editing in plant mitochondria and chloroplasts. Mol Biol Evol. 1997;14: 344–349. doi:10.1093/oxfordjournals.molbev.a025768

49. Lynch M. Streamlining and simplification of microbial genome architecture. Annu Rev Microbiol. 2006;60: 327–349. doi:10.1146/annurev.micro.60.080805.142300

50. Hammani K, Okuda K, Tanz SK, Chateigner-Boutin A-L, Shikanai T, Small I. A study of new Arabidopsis chloroplast RNA editing mutants reveals general features of editing factors and their target sites. Plant Cell. 2009;21: 3686–3699. doi:10.1105/tpc.109.071472

51. Sloan DB, MacQueen AH, Alverson AJ, Palmer JD, Taylor DR. Extensive Loss of RNA Editing Sites in Rapidly Evolving Silene Mitochondrial Genomes: Selection vs. Retroprocessing as the Driving Force. Genetics. 2010;185: 1369–1380. doi:10.1534/genetics.110.118000

52. Meers C, Keskin H, Storici F. DNA repair by RNA: Templated, or not templated, that is the question. DNA Repair. 2016;44: 17–21. doi:10.1016/j.dnarep.2016.05.002

53. Bader AS, Bushell M. iMUT-seq: high-resolution DSB-induced mutation profiling reveals prevalent homologous-recombination dependent mutagenesis. Nat Commun. 2023;14: 8419. doi:10.1038/s41467-023-44167-1

54. Lloyd AH, Rousseau-Gueutin M, Timmis JN, Sheppard AE, Ayliffe MA. Promiscuous Organellar DNA. In: Bock R, Knoop V, editors. Genomics of Chloroplasts and Mitochondria. Dordrecht: Springer Nature; 2012. pp. 201–221. doi:10.1007/978-94-007-2920-9_9

55. Wang D, Wu Y-W, Shih AC-C, Wu C-S, Wang Y-N, Chaw S-M. Transfer of chloroplast genomic DNA to mitochondrial genome occurred at least 300 Mya. Mol Biol Evol. 2007;24: 2040–2048. doi:10.1093/molbev/msm133

56. Adams KL, Daley DO, Qiu YL, Whelan J, Palmer JD. Repeated, recent and diverse transfers of a mitochondrial gene to the nucleus in flowering plants. Nature. 2000;408: 354–357. doi:10.1038/35042567

57. Wu Z, Sloan DB, Brown CW, Rosenblueth M, Palmer JD, Ong HC. Mitochondrial Retroprocessing Promoted Functional Transfers of rpl5 to the Nucleus in Grasses. Mol Biol Evol. 2017;34: 2340–2354. doi:10.1093/molbev/msx170

58. Suzuki JY, Sriraman P, Svab Z, Maliga P. Unique Architecture of the Plastid Ribosomal RNA Operon Promoter Recognized by the Multisubunit RNA Polymerase in Tobacco and Other Higher Plants. Plant Cell. 2003;15: 195–205. doi:10.1105/tpc.007914

59. Tiller N, Bock R. The Translational Apparatus of Plastids and Its Role in Plant Development. Mol Plant. 2014;7: 1105–1120. doi:10.1093/mp/ssu022

60. Palomar VM, Jaksich S, Fujii S, Kuciński J, Wierzbicki AT. High-resolution map of plastid-encoded RNA polymerase binding patterns demonstrates a major role of transcription in chloroplast gene expression. Plant J. 2022;111: 1139–1151. doi:10.1111/tpj.15882

61. Campo EM del, Sabater B, Martín M. Transcripts of the ndhH–D operon of barley plastids: possible role of unedited site III in splicing of the ndhA intron. Nucleic Acids Res. 2000;28: 1092–1098.

62. Serrot PH, Sabater B, Martín M. Expression of the ndhCKJ operon of barley and editing at the 13th base of the mRNA of the ndhC gene. Biol Plant. 2008;52: 347–350. doi:10.1007/s10535-008-0071-y

63. Nhat Nam N, Pham Anh Thi N, Do HDK. New Insights into the Diversity of Mitochondrial Plastid DNA. Genome Biol Evol. 2024;16: evae184. doi:10.1093/gbe/evae184

64. Wang X-C, Chen H, Yang D, Liu C. Diversity of mitochondrial plastid DNAs (MTPTs) in seed plants. Mitochondrial DNA Part DNA Mapp Seq Anal. 2018;29: 635–642. doi:10.1080/24701394.2017.1334772

65. Barbrook AC, Howe CJ, Purton S. Why are plastid genomes retained in non-photosynthetic organisms? Trends Plant Sci. 2006;11: 101–108. doi:10.1016/j.tplants.2005.12.004

66. Smith DR. Extending the Limited Transfer Window Hypothesis to Inter-organelle DNA Migration. Genome Biol Evol. 2011;3: 743–748. doi:10.1093/gbe/evr068

67. Golczyk H, Greiner S, Wanner G, Weihe A, Bock R, Börner T, et al. Chloroplast DNA in Mature and Senescing Leaves: A Reappraisal. Plant Cell. 2014;26: 847–854. doi:10.1105/tpc.113.117465

68. Fujie M, Kuroiwa H, Kawano S, Kuroiwa T. Studies on the behavior of organelles and their nucleoids in the root apical meristem of Arabidopsis thaliana (L.) Col. Planta. 1993;189: 443–452. doi:10.1007/BF00194444

69. Fujie M, Kuroiwa H, Suzuki T, Kawano S, Kuroiwa T. Organelle DNA Synthesis in the Quiescent centre of Arabidopsis thaliana (Col.). J Exp Bot. 1993;44: 689–693. doi:10.1093/jxb/44.4.689

70. Sloan DB, Alverson AJ, Chuckalovcak JP, Wu M, McCauley DE, Palmer JD, et al. Rapid evolution of enormous, multichromosomal genomes in flowering plant mitochondria with exceptionally high mutation rates. PLOS Biol. 2012;10: e1001241. doi:10.1371/journal.pbio.1001241

71. Lassalle F, Périan S, Bataillon T, Nesme X, Duret L, Daubin V. GC-content evolution in bacterial genomes: the biased gene conversion hypothesis expands. PLOS Genet. 2015;11: e1004941. doi:10.1371/journal.pgen.1004941

72. Thomas CM, Nielsen KM. Mechanisms of, and barriers to, horizontal gene transfer between bacteria. Nat Rev Microbiol. 2005;3: 711–721. doi:10.1038/nrmicro1234

73. Wu Z, Cuthbert JM, Taylor DR, Sloan DB. The massive mitochondrial genome of the angiosperm Silene noctiflora is evolving by gain or loss of entire chromosomes. Proc Natl Acad Sci. 2015;112: 10185–10191. doi:10.1073/pnas.1421397112

74. Zwonitzer KD, Tressel LG, Wu Z, Kan S, Broz AK, Mower JP, et al. Genome copy number predicts extreme evolutionary rate variation in plant mitochondrial DNA. Proc Natl Acad Sci. 2024;121: e2317240121. doi:10.1073/pnas.2317240121

75. Arias-Agudelo LM, González F, Isaza JP, Alzate JF, Pabón-Mora N. Plastome reduction and gene content in New World Pilostyles (Apodanthaceae) unveils high similarities to African and Australian congeners. Mol Phylogenet Evol. 2019;135: 193–202.

76. Molina J, Hazzouri KM, Nickrent D, Geisler M, Meyer RS, Pentony MM, et al. Possible loss of the chloroplast genome in the parasitic flowering plant Rafflesia lagascae (Rafflesiaceae). Mol Biol Evol. 2014;31: 793–803.

77. Su H-J, Barkman TJ, Hao W, Jones SS, Naumann J, Skippington E, et al. Novel genetic code and record-setting AT-richness in the highly reduced plastid genome of the holoparasitic plant Balanophora. Proc Natl Acad Sci. 2019;116: 934–943.

78. Parkinson CL, Mower JP, Qiu Y-L, Shirk AJ, Song K, Young ND, et al. Multiple major increases and decreases in mitochondrial substitution rates in the plant family Geraniaceae. BMC Evol Biol. 2005;5: 73. doi:10.1186/1471-2148-5-73

79. Miller-Messmer M, Kühn K, Bichara M, Le Ret M, Imbault P, Gualberto JM. RecA-dependent DNA repair results in increased heteroplasmy of the Arabidopsis mitochondrial genome. Plant Physiol. 2012;159: 211–226. doi:10.1104/pp.112.194720

80. Senkler J, Rugen N, Eubel H, Hegermann J, Braun H-P. Absence of Complex I implicates rearrangement of the respiratory chain in European mistletoe. Curr Biol. 2018;28: 1606–1613.e4. doi:10.1016/j.cub.2018.03.050

81. Jobson RW, Nielsen R, Laakkonen L, Wikström M, Albert VA. Adaptive evolution of cytochrome c oxidase: Infrastructure for a carnivorous plant radiation. Proc Natl Acad Sci U S A. 2004;101: 18064–18068. doi:10.1073/pnas.0408092101

82. Maclean AE, Hertle AP, Ligas J, Bock R, Balk J, Meyer EH. Absence of Complex I is associated with diminished respiratory chain function in European mistletoe. Curr Biol. 2018;28: 1614–1619.e3. doi:10.1016/j.cub.2018.03.036

83. Patiño S, Aalto T, Edwards AA, Grace J. Is Rafflesia an endothermic flower? New Phytol. 2002;154: 429–437.

84. Wang X, Morton JA, Pellicer J, Leitch IJ, Leitch AR. Genome downsizing after polyploidy: mechanisms, rates and selection pressures. Plant J. 2021;107: 1003–1015. doi:10.1111/tpj.15363

85. Ibarra-Laclette E, Lyons E, Hernández-Guzmán G, Pérez-Torres CA, Carretero-Paulet L, Chang T-H, et al. Architecture and evolution of a minute plant genome. Nature. 2013;498: 94–98. doi:10.1038/nature12132

86. Zedek F, Šmerda J, Halasová A, Adamec L, Veleba A, Plačková K, et al. The smallest angiosperm genomes may be the price for effective traps of bladderworts. Ann Bot. 2024; mcae107. doi:10.1093/aob/mcae107

87. Doyle J, Doyle JL. Genomic plant DNA preparation from fresh tissue-CTAB method. Phytochem Bull. 1987;19: 11–15.

88. Stewart CN, Via LE. A rapid CTAB DNA isolation technique useful for RAPD fingerprinting and other PCR applications. BioTechniques. 1993;14: 748–750.

89. Thieret JW. The genera of Orobanchaceae in the Southeastern United States. J Arnold Arbor. 1971;52: 404–434.

90. Krueger F. Trim Galore: a wrapper tool around Cutadapt and FastQC to consistently apply quality and adapter trimming to FastQ files, with some extra functionality for MspI-digested RRBS-type (Reduced Representation Bisufite-Seq) libraries. URL Httpwww Bioinforma Babraham Ac UkprojectstrimgaloreDate Access 28042016. 2012.

91. Jin J-J, Yu W-B, Yang J-B, Song Y, DePamphilis CW, Yi T-S, et al. GetOrganelle: a fast and versatile toolkit for accurate de novo assembly of organelle genomes. Genome Biol. 2020;21: 1–31.

92. Wick RR, Schultz MB, Zobel J, Holt KE. Bandage: interactive visualization of de novo genome assemblies. Bioinformatics. 2015;31: 3350–3352. doi:10.1093/bioinformatics/btv383

93. Langmead B, Salzberg SL. Fast gapped-read alignment with Bowtie 2. Nat Methods. 2012;9: 357–359.

94. Li H, Durbin R. Fast and accurate short read alignment with Burrows–Wheeler transform. Bioinformatics. 2009;25: 1754–1760. doi:10.1093/bioinformatics/btp324

95. Milne I, Bayer M, Cardle L, Shaw P, Stephen G, Wright F, et al. Tablet—next generation sequence assembly visualization. Bioinformatics. 2010;26: 401–402. doi:10.1093/bioinformatics/btp666

96. Heldt H-W, Piechulla B. A plant cell has three different genomes. In: Heldt H-W, Piechulla B, editors. Plant Biochemistry (Fifth Edition). Academic Press; 2021. pp. 469–501. doi:10.1016/B978-0-12-818631-2.00020-9

97. Marçais G, Kingsford C. A fast, lock-free approach for efficient parallel counting of occurrences of k-mers. Bioinforma Oxf Engl. 2011;27: 764–770.

98. Minh BQ, Schmidt HA, Chernomor O, Schrempf D, Woodhams MD, Von Haeseler A, et al. IQ-TREE 2: New models and efficient methods for phylogenetic inference in the genomic era. Mol Biol Evol. 2020;37: 1530–1534.

99. Smith SA, O’Meara BC. treePL: divergence time estimation using penalized likelihood for large phylogenies. Bioinformatics. 2012;28: 2689–2690.

100. Wynn EL, Christensen AC. Repeats of unusual size in plant mitochondrial genomes: identification, incidence and evolution. G3 GenesGenomesGenetics. 2019;9: 549–559. doi:10.1534/g3.118.200948

101. Quinlan AR, Hall IM. BEDTools: a flexible suite of utilities for comparing genomic features. Bioinforma Oxf Engl. 2010;26: 841–842.

102. Csardi G, Nepusz T. The igraph software package for complex network research. InterJournal. 2006;Complex Systems: 1695.

103. Marçais G, Delcher AL, Phillippy AM, Coston R, Salzberg SL, Zimin A. MUMmer4: A fast and versatile genome alignment system. PLOS Comput Biol. 2018;14: e1005944. doi:10.1371/journal.pcbi.1005944

104. Edera AA, Small I, Milone DH, Sanchez-Puerta MV. Deepred-Mt: Deep representation learning for predicting C-to-U RNA editing in plant mitochondria. Comput Biol Med. 2021;136: 104682. doi:10.1016/j.compbiomed.2021.104682

105. Hanazawa M, Narushima H, Minaka N. Generating most parsimonious reconstructions on a tree: A generalization of the Farris-Swofford-Maddison method. Discrete Appl Math. 1995;56: 245–265. doi:10.1016/0166-218X(94)00089-V

106. Paradis E, Schliep K. ape 5.0: an environment for modern phylogenetics and evolutionary analyses in R. Bioinformatics. 2019;35: 526–528. doi:10.1093/bioinformatics/bty633

107. Katoh K, Standley DM. MAFFT multiple sequence alignment software version 7: improvements in performance and usability. Mol Biol Evol. 2013;30: 772–780.

108. Ranwez V, Harispe S, Delsuc F, Douzery EJ. MACSE: Multiple Alignment of Coding SEquences accounting for frameshifts and stop codons. PLOS One. 2011;6: e22594.

109. Yang Z. PAML 4: phylogenetic analysis by maximum likelihood. Mol Biol Evol. 2007;24: 1586–1591.

110. Kosakovsky Pond SL, Poon AFY, Velazquez R, Weaver S, Hepler NL, Murrell B, et al. HyPhy 2.5—A customizable platform for evolutionary hypothesis testing using phylogenies. Mol Biol Evol. 2020;37: 295–299. doi:10.1093/molbev/msz197

111. Revell LJ. phytools: an R package for phylogenetic comparative biology (and other things). Methods Ecol Evol. 2012;3: 217–223. doi:10.1111/j.2041-210X.2011.00169.x

112. Orme D, Freckleton R, Thomas G, Petzoldt T, Fritz S, Isaac N, et al. caper: comparative analyses of phylogenetics and evolution in R. 2023. Available: https://cran.r-project.org/web/packages/caper/index.html

113. Rambaut A, Drummond AJ, Xie D, Baele G, Suchard MA. Posterior summarization in Bayesian phylogenetics using Tracer 1.7. Syst Biol. 2018;67: 901–904. doi:10.1093/sysbio/syy032

