## Supplementary Information for "Recombination and retroprocessing in broomrapes reveal a universal roadmap for mitochondrial evolution in heterotrophic plants"

**Supplementary figures**

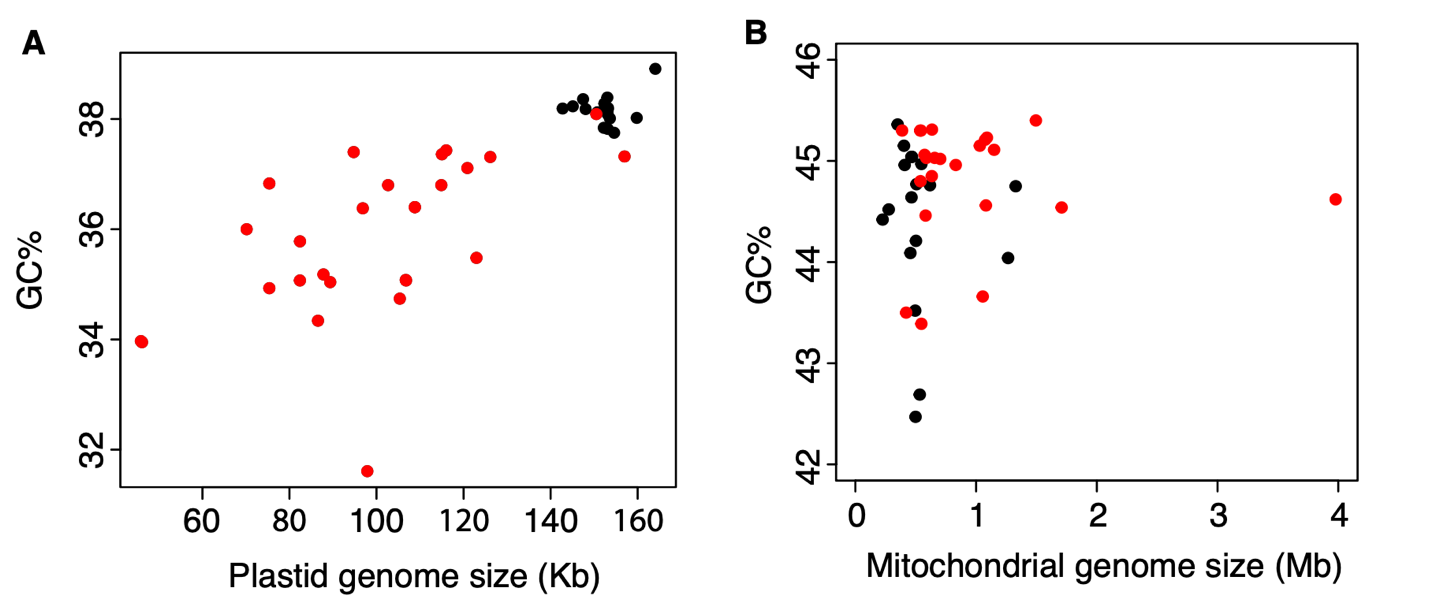

**Figure S1** Correlation between genome size and GC content in organellar genomes. Free-living and hemiparasitic species are represented by black dots and holoparasites are represented by red dots. (A) Plastid genome size shows a trend of positive correlation with GC content although not statistically significant (phylogenetic generalized least squares *p*-value = 0.085). (B) Mitochondrial genome size shows no correlation with GC content.

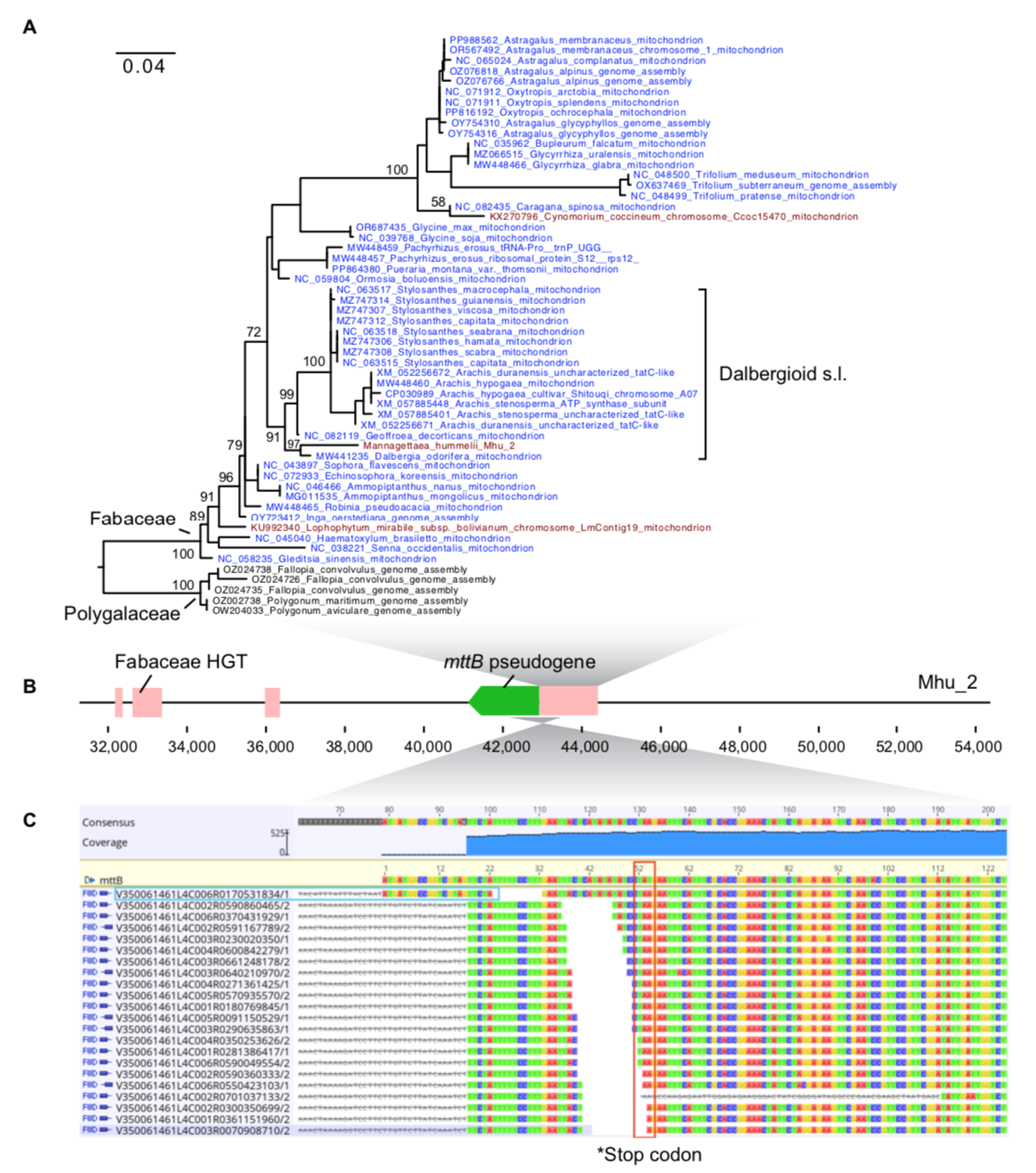

**Figure S2** Insertion of horizontally transferred (HGT) mitochondrial DNA from legume host leads to loss of function of the *mttB* gene in the parasitic *Mannagettaea hummelii*. (A) Maximum likelihood phylogeny of a 1,100 bp insertion flanking the 5’ end of *mttB* inferred from IQ-TREE. Branch lengths are in mutation units and branch support is calculated using 1000 ultrafast bootstrap replication. This segment is only found in Fabaceae (blue), Polygalaceae (black), and three parasitic plants (red). *Mannagettaea* is well nested within the Dalbergioid s.l. clade and clusters with *Dalbergia*. The other two parasitic species *Lophophytum mirabile* (Balanophoraceae) and *Cynomorium coccineum* (Cynomoriaceae) likely obtained this sequence via host-to-parasite HGT from Fabaceae as well*.* (B) Physical location of *mttB* and adjacent HGT from Fabaceae. (C) Reads mapping supports the premature stop codon at 5’ end of *mttB*. One read aligned to the intact *mttB* reference sequence, suggesting a potentially intact nuclear copy. Although our sequencing coverage is insufficient to recover a full-length nuclear copy of *mttB*.

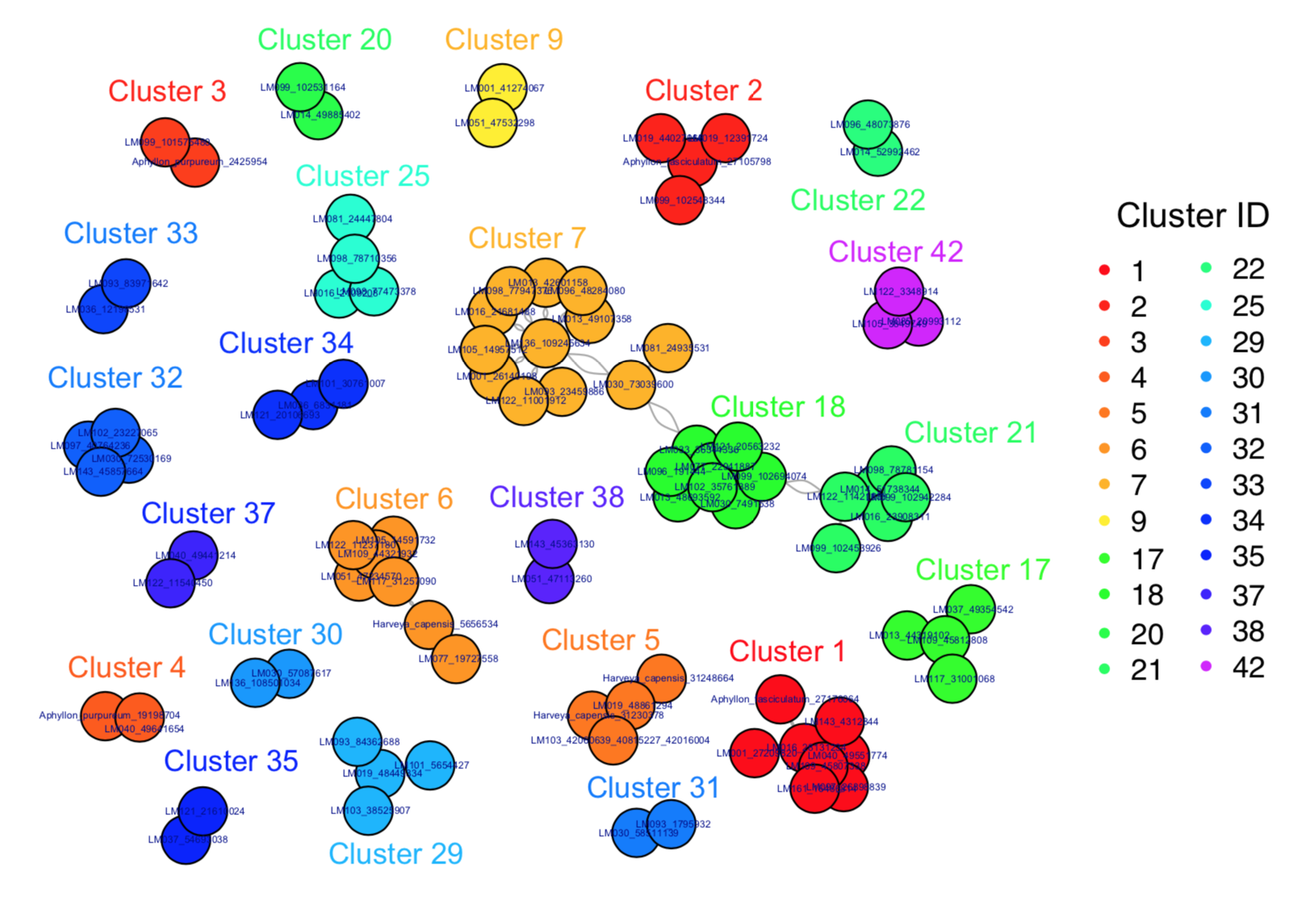

**Figure S3** Convergent evolution of dispersed repeats in Orobanchaceae mitogenomes. Structural repeats identified from De Bruijn graphs were clustered based on sequence similarity. Twenty-four clusters with at least two independent origins of the same repeats are listed. Each cluster is color coded by the cluster ID. Details regarding the length of the repeat unit, the length of homologous sequences, and sequence annotation are provided in Table S6.

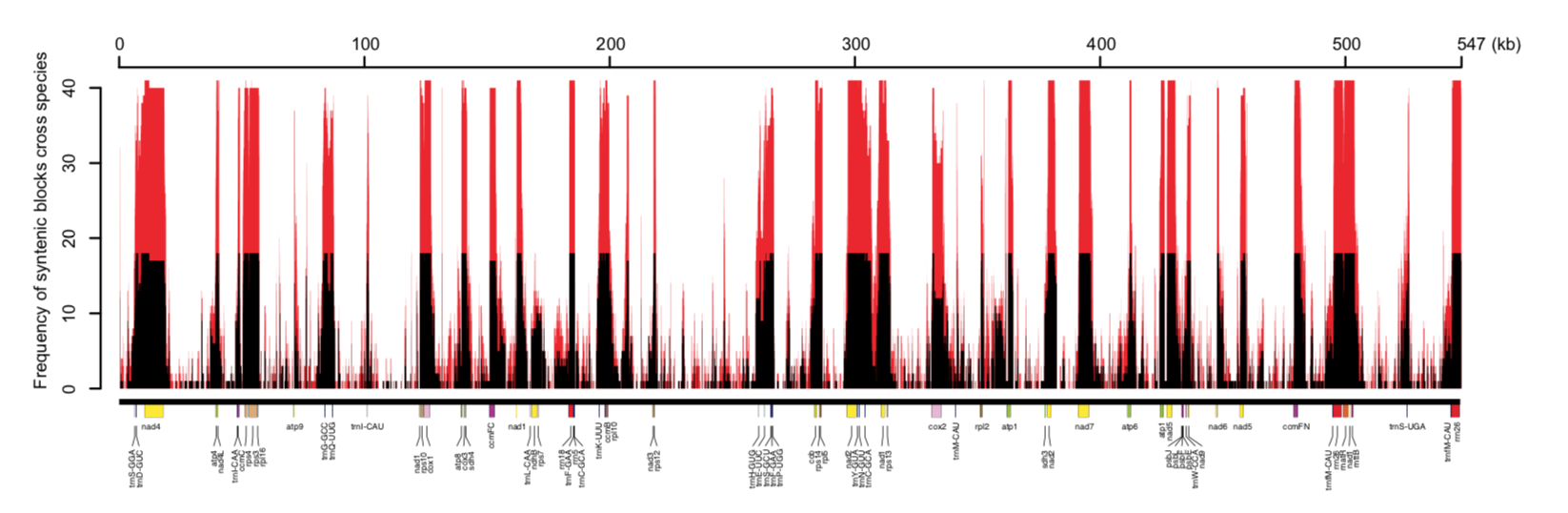

**Figure S4** Lack of synteny outside coding regions in the mitogenomes of Orobanchaceae. The mitogenome of *Rehmannia glutinosa* (x-axis) was aligned with each species using Mummer v4.0. The number of Orobanchaceae species in synteny with *Rehmannia* was enumerated for each site and visualized in the y-axis. Black bars represent hemiparasites and red bars represent holoparasites. Peaks represent hotspots of conserved syntenetic regions.

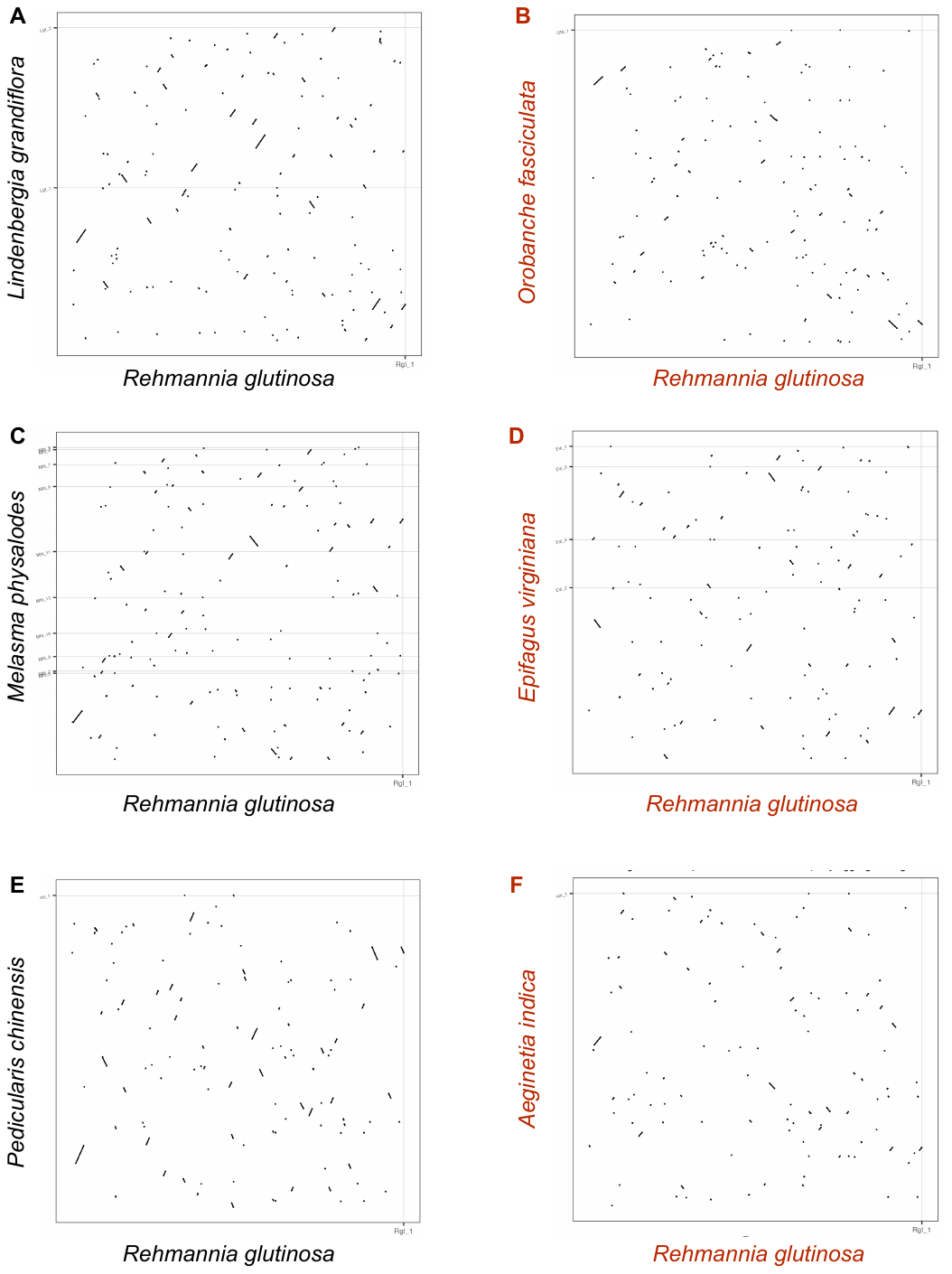

**Figure S5** Selected pairwise genome alignments with *Rehmannia glutinosa*. Mitogenomes from (A) *Lindenbergia grandiflora*, (B) *Aphyllon fasciculatum*, (C) *Melasma physalodes*, (D) *Epifagus virginiana*, (E) *Pedicularis chinensis*, and (F) *Aeginetia indica* are aligned to *Rehmannia* using Mummer v4.0. Aligned segments are represented as dots and lines and holoparasitic species are highlighted in red.

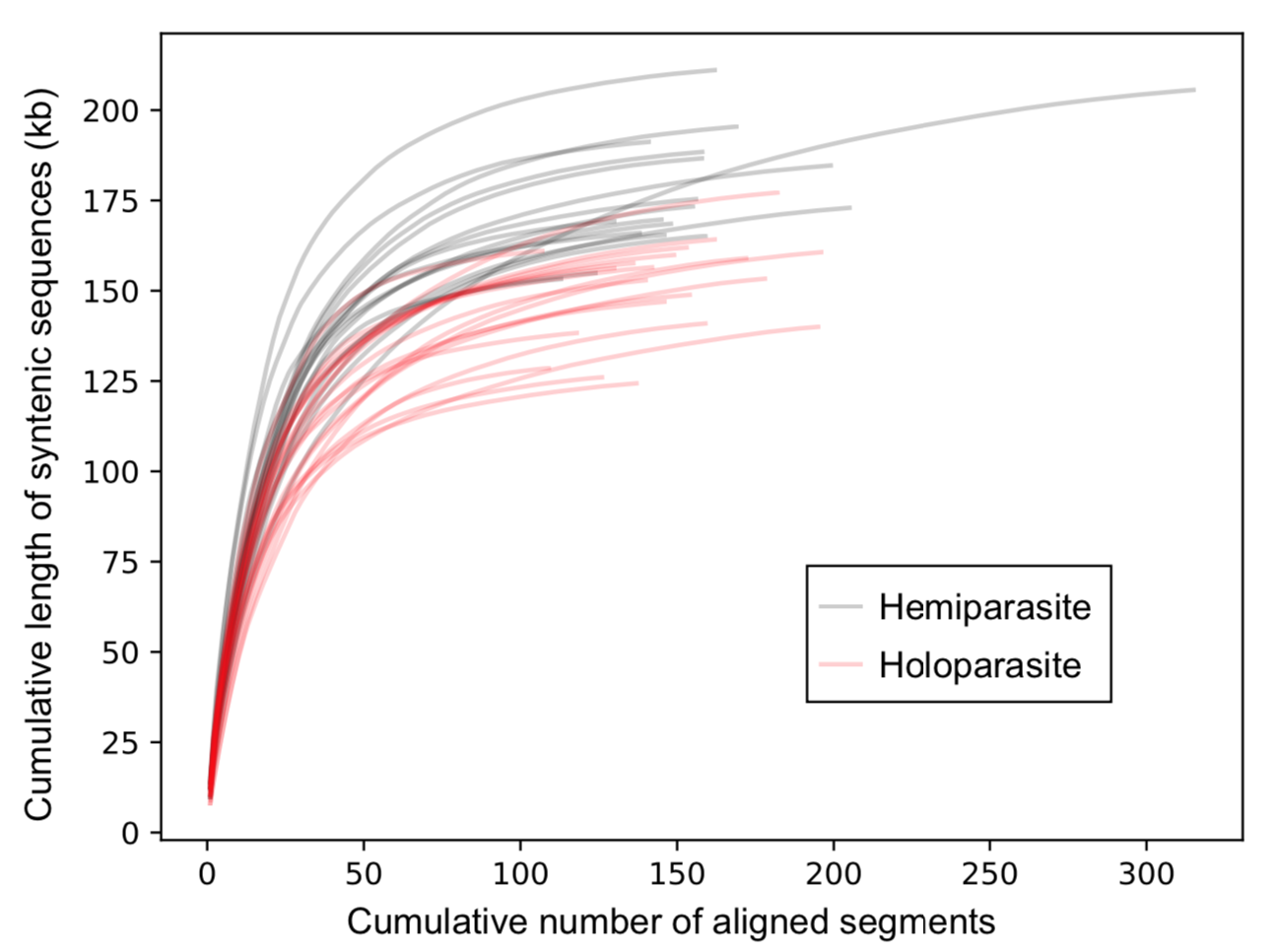

**Figure S6** Cumulative length of syntenic mitogenome regions in hemi- and holo-parasites. The mitogenome of each species was aligned to *Rehmannia glutinosa* using Mummer. The aligned segments were ordered by length, and the cumulative length of all segments was plotted. The total length and number of syntenic regions define the end point of each curve, while the initial slope of each curve reflects the proportion of longer blocks. Gray lines represent hemiparasites and red lines represent holoparasites.

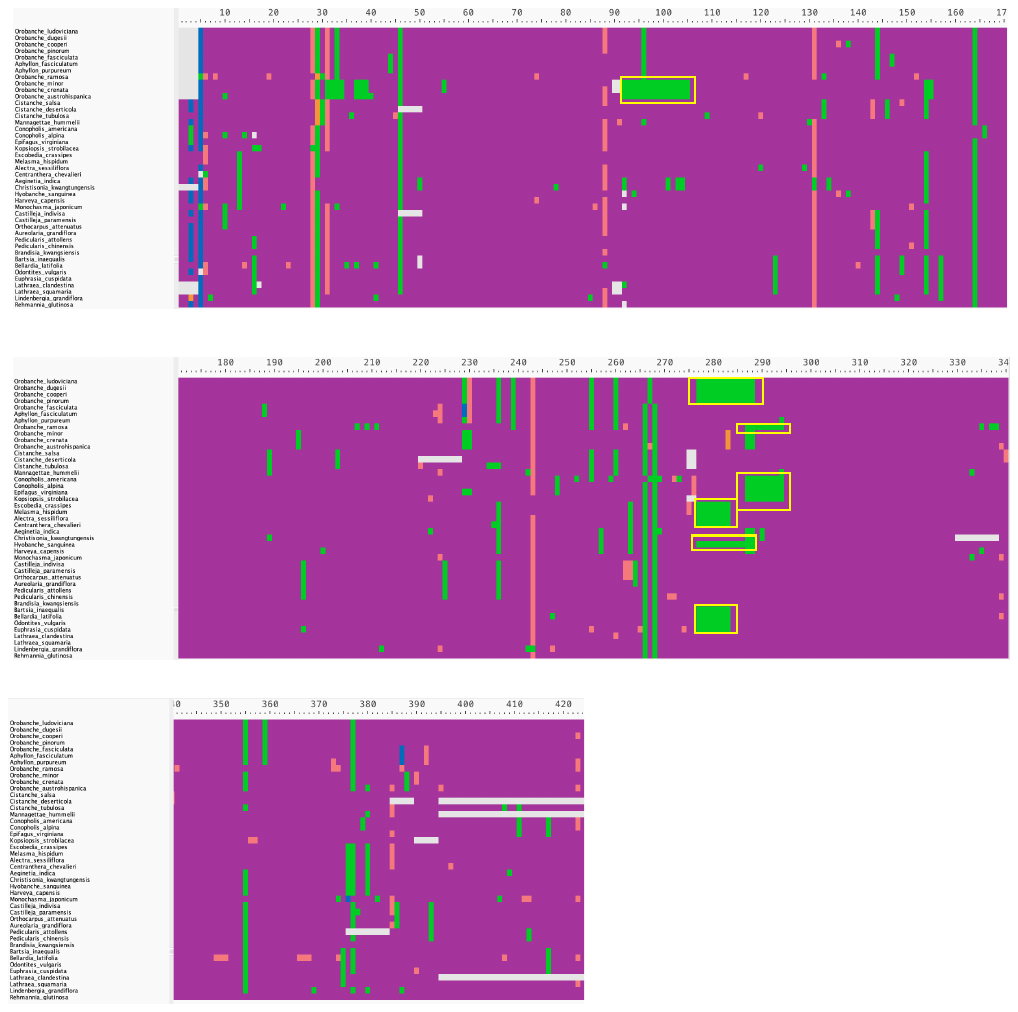

**Figure S7** Sequence alignment of predicted C-to-U RNA editing sites for 27 core mitochondrial genes (defined in Table S7). RNA editing is predicted by Deepred-Mt and colored in purple. Only sites predicted with editing sites in more than 2 species are shown. Green cells represent thymine and hence the loss of RNA editing; blue cells represent adenine; red cells represent cytosine. Continuous stretches of thymine highlighted with yellow rectangles are likely results of gene conversion of reverse-transcribed mRNAs.

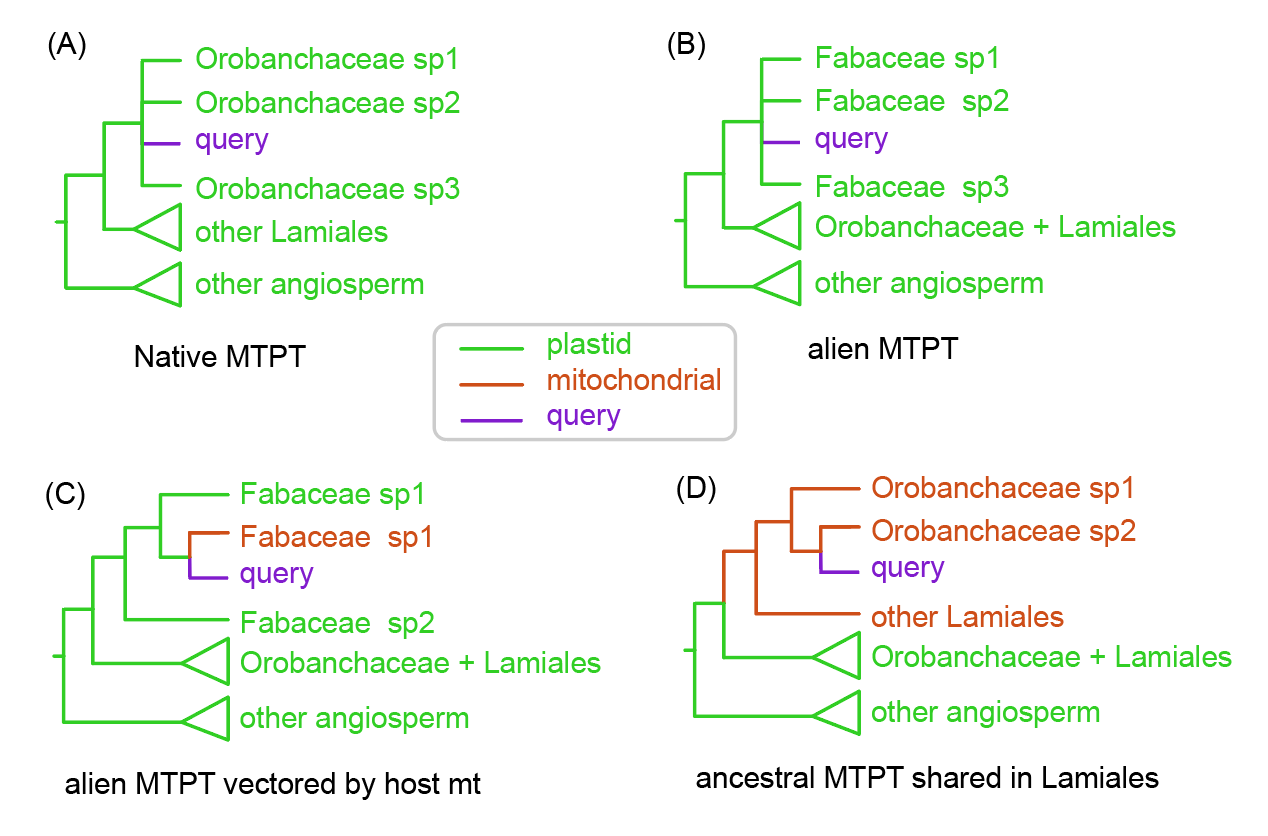

**Figure S8** Classification of mitochondrial plastid DNA (MTPT) based on the genomic location and taxonomy of the donor. Four common types of MTPT are illustrated with their expected phylogeny. The query sequence is shown in purple, the plastid sequence is shown in green, and the mitochondrial sequence in shown in brown.

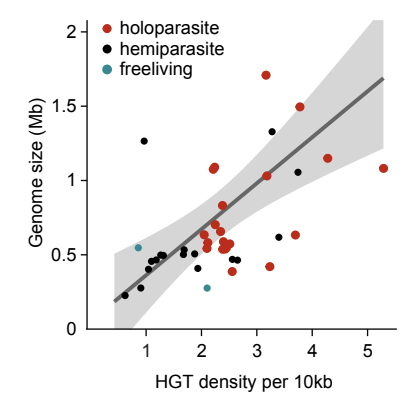

**Figure S9** Positive correlation between mitochondrial genome size and density of horizontal gene transfer (HGT) in Orobanchaceae. HGT density is defined as the number of HGTs per 10 kb. Species are color coded according to lifestyle.

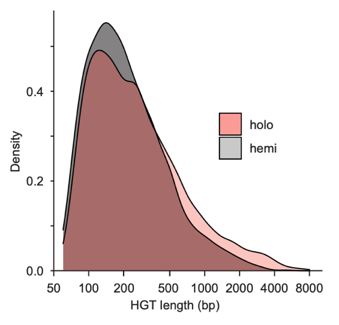

**Figure S10** Distribution of HGT segment size. HGTs from holoparasites are colored in red. The x-axis of HGT length in base pair (bp) is log-transformed to aid with visualization.

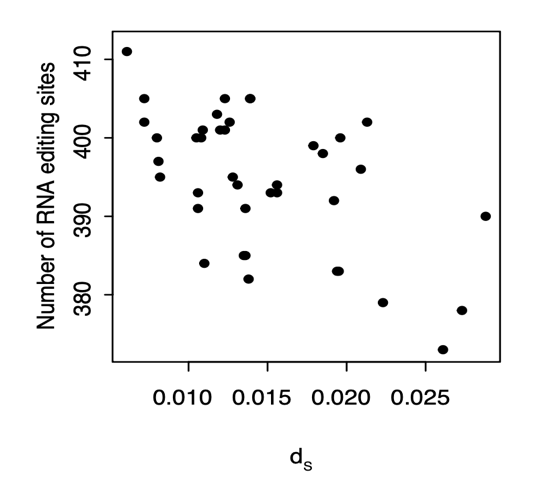

**Figure S11** Negative correlation between synonymous substitution rates (*d_S_*) and RNA editing sites in Orobanchaceae (phylogenetic generalized least squares *p*-value = 8.6e-4).

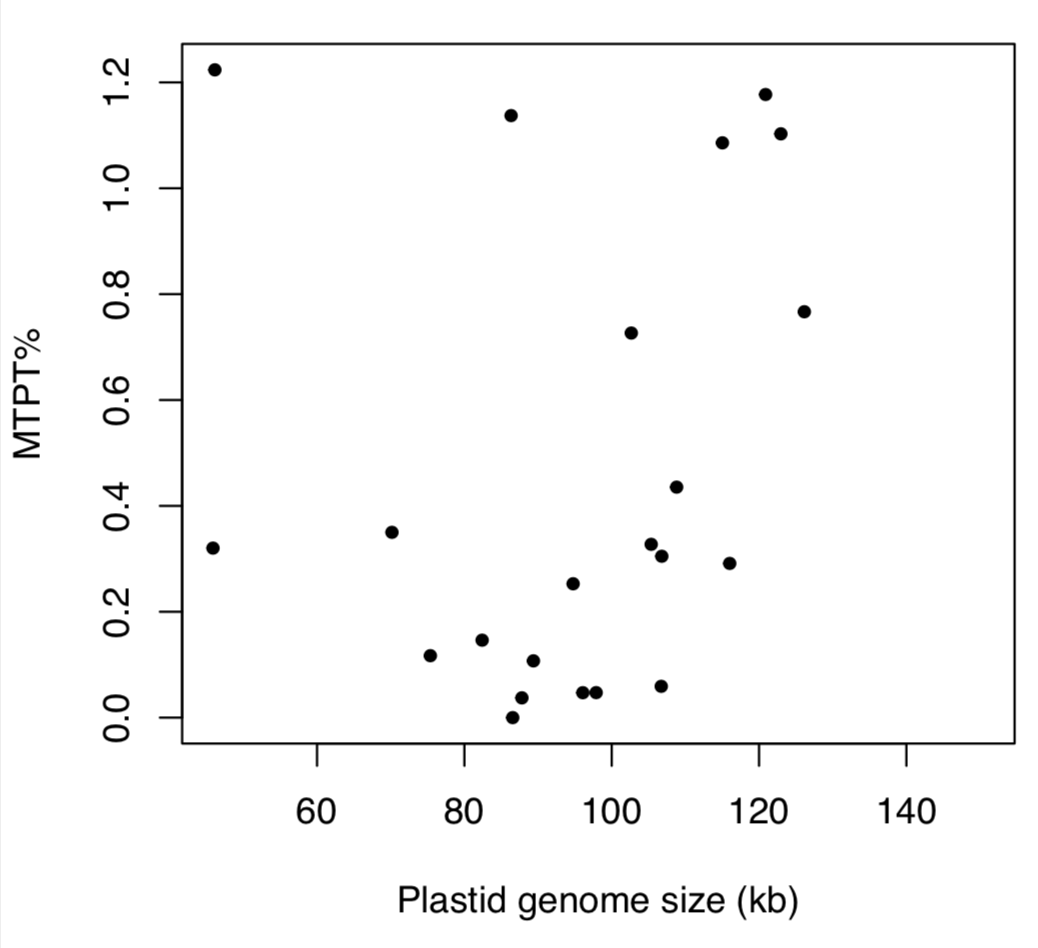

**Figure S12** Lack of correlation between plastid genome size and mitochondrial plastid DNA (MTPT) in holoparasitic Orobanchaceae. Phylogenetic generalized least squares test *p*-value = 0.45, adjusted R^2^ = -0.04753.

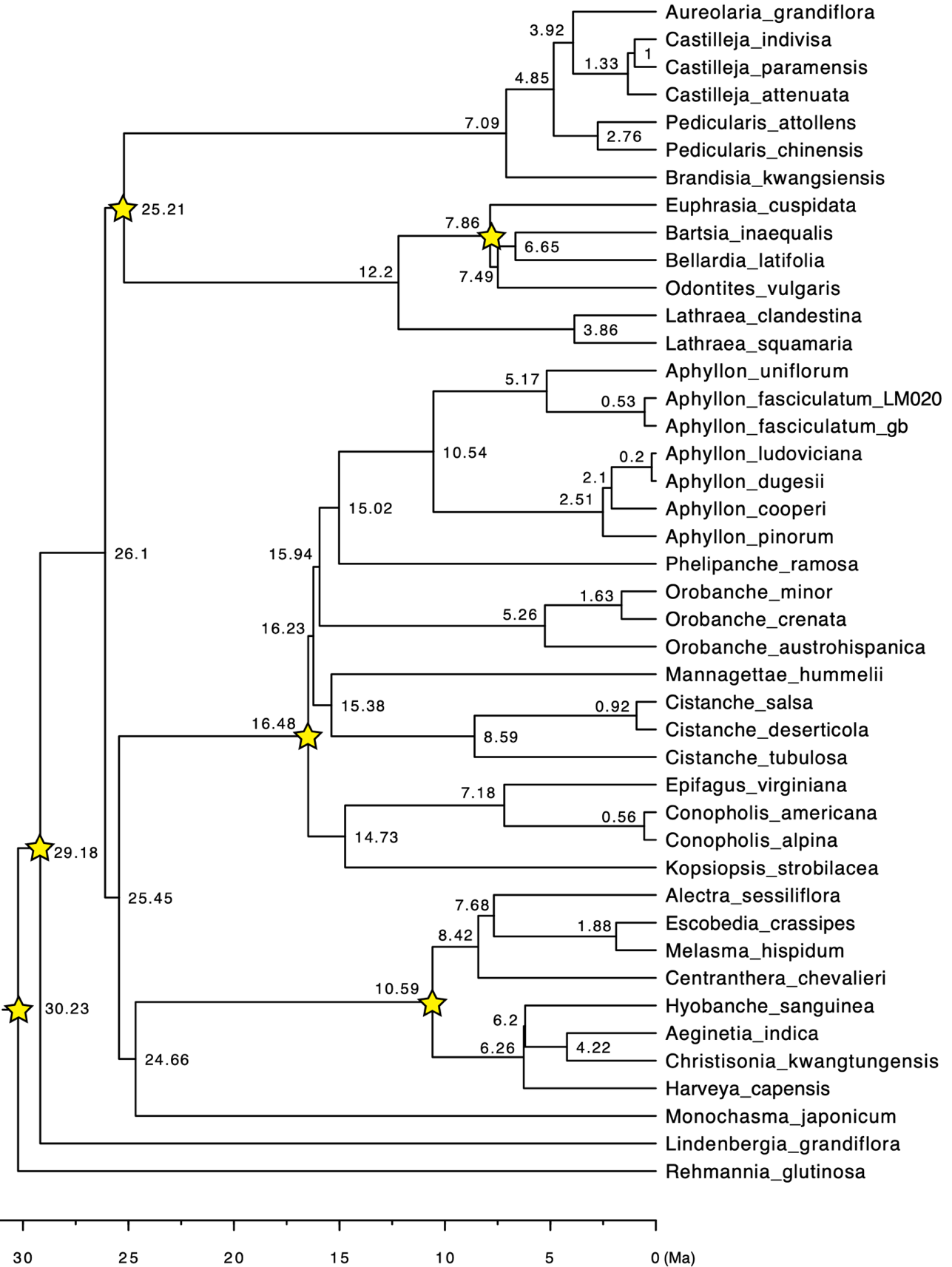

**Figure S13** Time tree of Orobanchaceae inferred from the penalized likelihood method implemented in TreePL v1.0. The mitochondrial phylogeny from Fig. 1 was used as input and the six secondary calibrations from Mortimer et al. (2022) were shown as stars on the nodes. The node age was shown in millions of years ago (Ma).

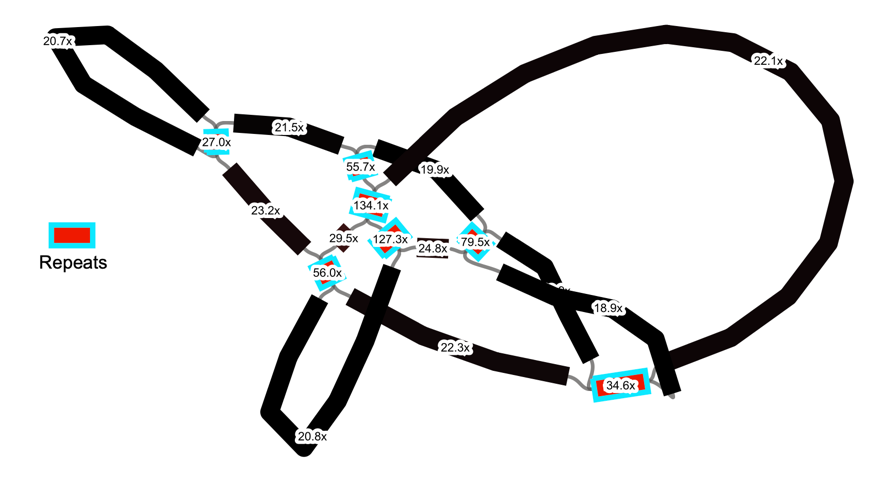

**Figure S14** Repeat identification using De Bruijn assembly graphs. The kmer coverage for each node was labeled on the example De Bruijn graph. Nodes with three or more connections and higher kmer coverage are identified as repeats.

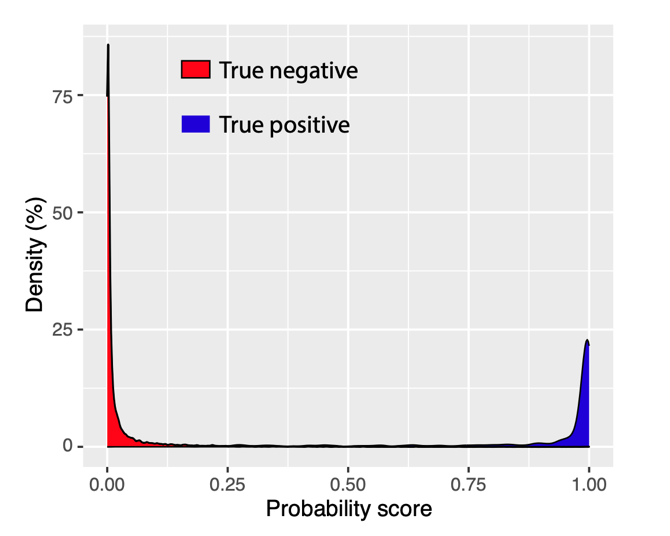

**Figure S15** Distribution of probability score of RNA editing sites predicted by Deepred-mt in experimentally verified sequences with RNA editing. Scores for the true positive sites are colored in blue and scores for the true negative sites are colored in red.

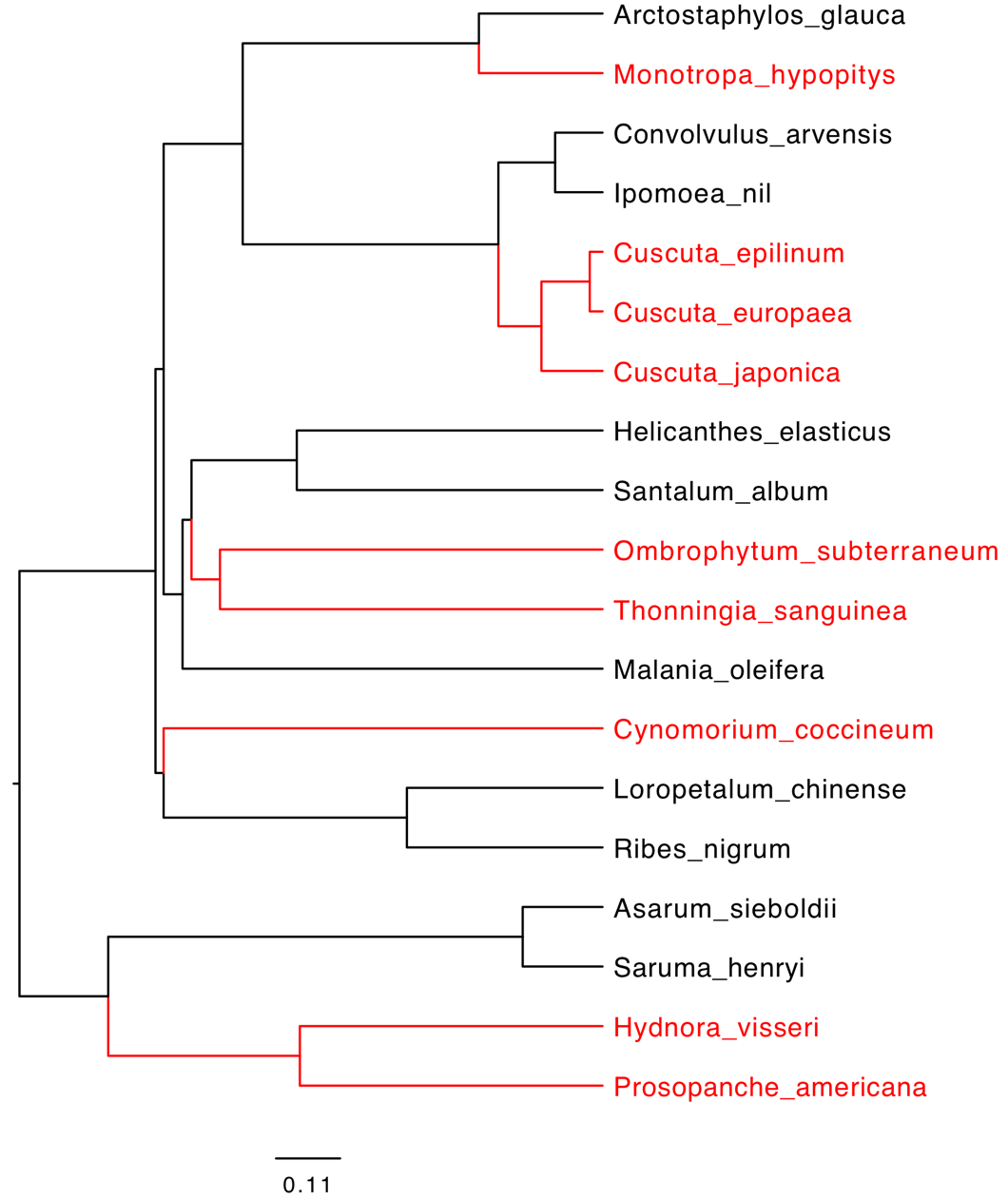

**Figure S16** Ultrametric tree of additional non-photosynthetic angiosperms inferred from the penalized likelihood method. Non-photosynthetic species are highlighted in red. A plastid phylogeny inferred from 24 core mitochondrial genes listed in Table S7 was used to generate an ultrametric tree using the chronos function from the R package ape.

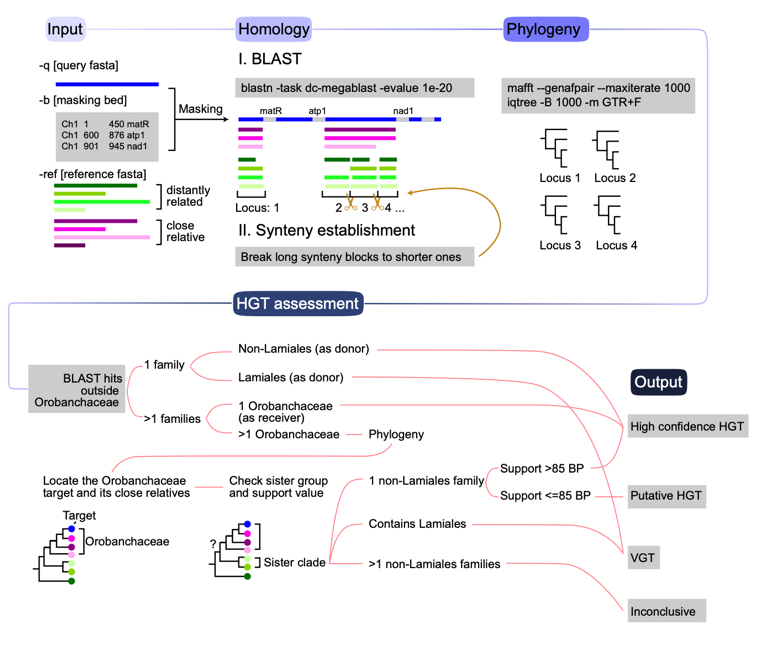

**Figure S17** Bioinformatic pipeline of HGTScanner to identify potential horizontal gene transfer in mitochondrial genomes. For a detailed description of the pipeline, see Supplementary Note 2.

**Supplementary tables**

**Table S1** Taxon sampling and voucher information for 45 Orobanchaceae species sampled in this study. The DNA extraction ID, GenBank SRA accession number, herbarium voucher information, and Illumina data size are provided for newly generated sequencing data. The GenBank ID and voucher information (when available) are provided for public data. An em dash represents “not applicable”.

**Table S2** Summary of plastid genome assembly in Orobanchaceae. GenBank accession numbers of plastid genome assemblies are provided for newly generated (labeled by ‘new’) and published (labeled by ‘published’) accessions. The plastid genomes of 45 Orobanchaceae species are all circularized. The total plastid genome size, GC content, size of the inverted repeat (IR) region, short single copy region (SSC), and long single copy region (LSC) are calculated. Unusual structural modifications are recorded as ‘Note’. An em dash represents “not applicable”.

**Table S3** Plastid, mitochondrial, and nuclear genome base coverage. Only species sequenced by us were used because of our unbiased, non-enrichment library preparation procedure. Base coverage of the plastid and mitochondrial assembly is estimated by reads mapping using BWA-MEM and Bowtie. The results reported in the main text are derived from BWA-MEM only. Base coverage of the nuclear genome is estimated using the kmer distribution with a kmer size of 21 in Jellyfish v2.3.0.

**Table S4** Summary of mitochondrial genome assembly. GenBank accession numbers of mitochondrial genome assemblies are provided for newly generated (labeled by ‘new’) and published (labeled by ‘published’) accessions. Based on the quality of the De Bruijn assembly graph from GetOrganelle, mitochondrial genome assemblies are classified as ‘Circularized’, ‘Complete draft’ (enclosed graph), ‘Nearly complete draft’ (open graph), or ‘Failed’ (no assembly produced). We also quantified the number of assembly scaffolds, GC content, genome size, MTPT (mitochondrial plastid DNA) content, repeat content, horizontal gene transfer (HGT) content, coding region length, and length of sequences syntenic to *Rehmannia* *glutinosa* for each species. An em dash represents “not applicable”. Assembly errors in the published mitochondrial genomes identified by reads mapping are recorded in the ‘Note’.

**Table S5** Mitochondrial gene and intron content in Orobanchaceae. For genes, “1” stands for intact sequences, “ψ” stands for pseudogenized relic sequences, “lost” stands for completely undetectable sequences, and “HGT” stands for potential horizontal gene transfers. Introns are highlighted by gray cells and “c” stands for cis-spliced intron, “t” stands for trans-spliced intron, “ψ” stands for loss of intron, and “HGT” stands for potential horizontal gene transfers.

**Table S6** Convergent evolution of dispersed repeats in Orobanchaceae mitochondrial genomes. Structural repeats identified from De Bruijn graphs were clustered based on sequence similarity using BLASTN and an e-value threshold of 1e-40. Twenty-four clusters with at least two independent origins of the same repeats are listed. Within each cluster, the ID and length of the repeat unit, the length of homologous sequences, and sequence annotation were listed.

**Table S7** Number of RNA editing sites per gene predicted by Deepred-Mt for 24 core mitochondrial protein coding genes.

**Table S8** Universally reduced mitochondrial plastid DNA (MTPT) in non-photosynthetic plants. MTPT content was calculated for five independently evolved non-photosynthetic lineages in Santalales, Piperales, Saxifragales, Solanales, and Ericales. For each species, the mitochondrial genome was compared to the plastid genome sequence of a reference species using BLASTN and an e-value threshold of 1e-70. The GenBank ID of the mitochondrial and plastid genomes used to calculate MTPT is provided for 19 species.

**Table S9** Donor and receiver lineages of alien mitochondrial plastid DNAs (MTPT) identified by the HGTScanner pipeline. Only high confidence alien MTPTs supported by phylogenies with >85 ultrafast bootstrap (UFBP) values are listed. The scaffold ID, position, type, donor, and UFBP of each MTPT are provided. Donors of these alien MTPTs may be plastid sequences (‘HGT from donor pt’), or MTPTs imbedded in donors’ mitochondrial genomes (‘HGT from donor mt’). The alignments and phylogenies supporting the identification of donor families and genera are provided in Supplementary Data S1. The names of the alignment data is coded as [Receiver scaffold ID]_[ Locus ID].fas and the phylogeny file is similarly coded as .

**Table S10** Lifestyle effects on selection in functional gene complexes based on the RELAX analyses. Mitochondrial genes were grouped based on their function. For each functional group, three branch models were tested, each using (i) tribe Orobancheae, (ii) three holoparasitic clades, and (iii) all parasitic Orobanchaceae as foreground branches, respectively. The log likelihood (LnL); corrected Akaike information criterion (AICc); *p*-value for differentiated selection in foreground lineages; selection intensity parameter *k* in RELAX; and likelihood ratio (LRT) from a likelihood ratio test of the RELAX null model and the RELAX alternative model were reported. A significant *p*-value (<0.05) and *k* < 1 indicates relaxed selection in the foreground branches; *k* > 1 indicates intensified selection.

**Table S11** Mitochondrial coding (CDS) and intergenic regions used for phylogenetic inference.

**Table 12** Universally conserved plastid genes used for phylogenetic inference.

**Table S13** Reference plastid genomes used to identify mitochondrial plastid DNA in Orobanchaceae. The family, species, and GenBank accessions are listed for 13 species broadly sampled across angiosperms with a focus on the known host lineages of Orobanchaceae.

**Supplementary Data**

Reviewer link: ###

Supplementary data supporting the findings of this study are available at FigShare (####). A brief description of the data is provided below.

- Species_ID.xlsx

The species names and IDs for the 45 species included in this study.

- 1_pt_assembly_annotation
  GetOrganelle assemblies and Geneious annotation of plastid genomes from 45 Orobanchaceae species
- 2_mt_assembly_annotation
  GetOrganelle assemblies and Geneious annotation of plastid genomes from 45 Orobanchaceae species
- 3_phylogeny
  Concatenated mitochondrial and plastid sequences IQ-TREE phylogenies inferred from these sequence data
- 4_rna_editing
  C-to-U RNA editing sites predicted by DeepRed-MT for 24 mitochondrial core genes
- 5_mtpt
  Sequence alignments and phylogenies supporting the classification of 1,094 mitochondrial plastid DNA (MTPT)
- 6_hgt
  Sequence alignments and phylogenies supporting the classification of mitochondrial horizontal gene transfer
- 7_molecular_evolution
  Codon alignments and program output of CODEML and HYPHY analyses

**Supplementary Note 1**

**Phylogenetic insights into Orobanchaceae**

The utility of mitochondrial loci as phylogenetic markers has been increasingly recognized in plant systematics [1]. Especially in heterotrophic plants, mitochondria offer slow-evolving, easily alignable sequences whereas the plastid DNA exhibits substantial gene loss and high substitution rates [2,3]. In Orobanchaceae, the comparison between mitochondrial and plastid phylogenies offers several important insights into the early diversification of Orobanchaceae.

First, the holoparasitic tribe Orobancheae was consistently placed as sister to tribe Buchnereae in both mitochondrial and plastid phylogeny (Fig. 1 in the main text). This relationship is also recovered in a recent study using more than nine hundred low-copy genes [4], but is in direct contrast with most previous studies where tribes Lindenbergieae, Cymbarieae, and Orobancheae form successive sisters to remaining Orobanchaceae [5–8]. Second, strongly conflicting placements are found in *Brandisia* and *Monochasma,* which allude to their hybrid origin. *Brandisia* is a member of the phylogenetically unstable East Asian *Pterygiella* clade . Our plastid phylogeny places *Brandisia* sister to tribe Rhinantheae whereas the mitochondrial phylogeny places it with Pedicularideae. The high support values, typical GC content, and moderate branch length of *Brandisia* rule out systematic biases for generating discrepancies. Instead, biological processes such as hybridization involving organelle capture are more likely, with Rhinantheae and Pedicularideae being the parental lineages. Future studies with more comprehensive taxon and gene sampling will be able to test the hybrid origin of *Brandisia* and *Monochasma* and evaluate whether these events impacted the entire *Pterygiella* and Cymbarieae clades they each represent. Third, the paraphyletic placements of the two *Aphyllon fasciculatum* accessions were also reported by [11] along with their distinct host preference for *Artemisia* (Asteraceae) and *Galium* (Rubiaceae) respectively. This example highlights the potentially widespread phenomenon of cryptic species complexes in parasitic Orobanchaceae characterized by host-driven speciation. In summary, mitochondria demonstrate significant utility in resolving and understanding the cause of controversial relationships at both deep and shallow levels.

**Supplementary Note 2**

**HGTScanner: a new bioinformatic tool to identify horizontal gene transfer in organellar genomes**

HGTScanner is a bioinformatic wrapper that establishes genome homology and identifies horizontal gene transfer (HGT) in plant mitochondrial genomes. The current version HGTScanner v1.0 contains two separate scripts HGTScanner_mtpt.py and HGTscanner_mt.py for HGT that originated from plastid and mitochondrial genomes, respectively. These two types of HGTs are treated differently because (i) the smaller size of the NCBI mitochondrial sequence collection makes it feasible and more efficient to conduct local BLAST searches while the huge size of the plastid sequence database makes it computationally highly inefficient to do so; (ii) the substantial mitochondrial genome structural variation challenges homology identification across difference species whereas the conserved plastome makes this relatively easy.

**Identifying alien MTPT with HGTScanner_mtpt**

To identify potential mitochondrial plastid DNAs (MTPTs) using HGTScanner_mtpt.py, we applied a two-step process by comparing the query sequence first to a custom database to identify potential HGTs, then to the entire NCBI GenBank database to finely place candidate HGTs. To accomplish this, we used a custom BLAST database containing plastid genomes from all 45 Orobanchaceae species sampled in this study (Table S3) and 13 additional angiosperms from Poaceae to Asteraceae (Table S13) to facilitate the identification of both native and alien plastid DNA. Homologous plastid regions in the mitochondrial genome were identified by BLASTN with a threshold of 1e-70 [12]. Homologous regions from the query sequence, Orobanchaceae plastids, and other angiosperm plastids were aligned with the MAFFT-einsi algorithm (--genapair --maxiterate 1000) [13]. A phylogeny was inferred using the optimum substitution model determined by ModelFinder in IQ-TREE v2.2.2.7 [14]. Branch support was evaluated using 1000 ultrafast bootstrap replicates (UFBP). Once the phylogenies were completed, the closest sister lineage of the query sequence was reported. If the query sequence was nested within Orobanchaceae, we reported this locus as a native MTPT. Otherwise, it was recognized as a potential alien MTPT and proceeded to the next step for verification. A total of 368 potential alien MTPTs were identified and processed in the next step.

To finely identify the donors of potential alien MTPTs, we compared them to the NCBI GenBank nucleotide (NT) database (accessed Nov 10, 2023 on https://blast.ncbi.nlm.nih.gov/Blast.cgi) using BLASTN. We also applied an e-value threshold of 1e-70 and restricted the results to the 100 top BLAST hits. The results were downloaded as xml file and converted to fasta sequences using the custom script ‘xml2seq.py’ (available on GitHub, <https://github.com/lmcai/Orobanchaceae_comparative_mitome/tree/main/5_mtpt>). These sequences were realigned with MAFFT and an IQ-TREE phylogeny was estimated for each locus using the same parameter settings described above. The sister clade and the branch support for the placement of the query sequence were reported in a summary spreadsheet.

Finally, to classify MTPT based on their evolutionary history, we manually examined the IQ-TREE phylogenies of the 368 potential alien MTPTs. Manual examination helps the identification of complex gene evolutionary histories such as: host plastid -> host mitochondrion -> parasite mitochondrion. These loci were classified into four categories: ‘native MTPT (V)’, ‘horizontally transferred MTPT from the host plastid (H)’, ‘horizontally transferred MTPT from the host mitochondria (H_mt)’, ‘MTPT shared by all Lamiales (A)’, and ‘inconclusive (U)’. Each of these scenarios left a unique phylogenetic signal:

I. Native MTPT (V)
   The query sequence is expected to nest within a clade consisted of Orobanchaceae plastid sequences (Fig. S8A).
II. Horizontally transferred MTPT from the host plastid (H)
    The query sequence is expected to nest within a clade consisted of non-Lamiales plastid sequences (Fig. S8B).
III. Horizontally transferred MTPT from the host mitochondria (H_mt)
     The query sequence is expected to cluster most closely with mitochondrial sequences from non-Lamiales species, then these sequences are nested within plastid sequences from the same non-Lamiales species (Fig. S8C).
IV. MTPT shared by all Lamiales (A)
    The query sequence is expected to cluster with other Lamiales mitochondrial sequences that nested within a clade consisted of Lamiales plastid sequences (Fig. S8D).
V. Inconclusive (U)

Other more complex scenarios.

**Identifying mitochondrial HGT with HGTScanner_mt**

To avoid excessive BLAST hits in the gene and MTPT regions, HGTscanner_mt relied on a user-supplied bed file to mask exons and MTPT in the query mitochondrial assembly. The masked query sequence was then BLAST against the entire Viridiplantae mitochondrial sequence database from NCBI GenBank (70,706 records; accessed 10 November 2023) using an e-value threshold of 1e-20. These BLAST hits were then ordered based on location and consolidated into longer synteny blocks using the ‘bedtools merge’ command from the pybedtools Python module. Each synteny block was further divided if more than 50% of the BLAST hits consisted of multiple genomic regions (Fig. S17). Such finer division is essential for the identification of shorter HGTs nested within long synteny blocks. It is also necessary due to the rapid breakdown of genomic synteny especially among distantly related species. Sequences from each synteny block were subsequently aligned using the MAFFT-einsi algorithm [13]. A maximum likelihood phylogeny was inferred for each block by IQ-TREE with 1000 ultrafast bootstrap replicates as described above for MTPT.

HGT was evaluated based on BLAST and phylogenetic evidence. The following criteria were used for classifying the query into four categories: ‘high confidence HGT’, ‘putative HGT’, ‘VGT’ (vertical gene transfer), and ‘inconclusive’ (Fig. S17).

I. High confidence HGT
   (1) Homology is found in Orobanchaceae and one non-Lamiales family (BLAST-based evidence); (2) homology is found in only one Orobanchaceae species and other land plant families (BLAST-based evidence); (3) the target Orobanchaceae parasite is nested well within a non-Lamiales family with >85 UFBP support (phylogeny-based evidence).
II. Putative HGT
    (1) The sister group of the target Orobanchaceae parasite contains only one non-Lamiales family, but the branch support is < 85 UFBP (phylogeny-based evidence).
III. VGT
     (1) Homology is found in Orobanchaceae and other Lamiales families (BLAST-based evidence); (2) the sister clade of the target Orobanchaceae species contains at least one Lamiales (phylogeny-based evidence).
IV. Inconclusive
    (1) The sister group of the target Orobanchaceae parasite contains more than one non-Lamiales family and thus cannot assign the donor of HGT confidently.

These stringent criteria are effective in removing false HGT due to phylogenetic uncertainty but may suffer from high false negative rates because most HGTs within Lamiales, especially from viable hosts in Lamiaceae and Plantaginaceae, will be classified as VGT.

**Reference**

1.	Lin Q, Braukmann TWA, Soto Gomez M, Mayer JLS, Pinheiro F, Merckx VSFT, et al. Mitochondrial genomic data are effective at placing mycoheterotrophic lineages in plant phylogeny. New Phytologist. 2022;236: 1908–1921. doi:10.1111/nph.18335

2.	Lam VKY, Darby H, Merckx VSFT, Lim G, Yukawa T, Neubig KM, et al. Phylogenomic inference in extremis: A case study with mycoheterotroph plastomes. Am J Bot. 2018;105: 480–494. doi:10.1002/ajb2.1070

3.	Naumann J, Der JP, Wafula EK, Jones SS, Wagner ST, Honaas LA, et al. Detecting and Characterizing the Highly Divergent Plastid Genome of the Nonphotosynthetic Parasitic Plant Hydnora visseri (Hydnoraceae). Genome Biol Evol. 2016;8: 345–363. doi:10.1093/gbe/evv256

4.	Xu Y, Zhang J, Ma C, Lei Y, Shen G, Jin J, et al. Comparative genomics of orobanchaceous species with different parasitic lifestyles reveals the origin and stepwise evolution of plant parasitism. Mol Plant. 2022;15: 1384–1399.

5.	Bennett JR, Mathews S. Phylogeny of the parasitic plant family Orobanchaceae inferred from phytochrome A. American Journal of Botany. 2006;93: 1039–1051.

6.	Fu W, Liu X, Zhang N, Song Z, Zhang W, Yang J, et al. Testing the Hypothesis of Multiple Origins of Holoparasitism in Orobanchaceae: Phylogenetic Evidence from the Last Two Unplaced Holoparasitic Genera, Gleadovia and Phacellanthus. Front Plant Sci. 2017;8. doi:10.3389/fpls.2017.01380

7.	Li X, Feng T, Randle C, Schneeweiss GM. Phylogenetic Relationships in Orobanchaceae Inferred From Low-Copy Nuclear Genes: Consolidation of Major Clades and Identification of a Novel Position of the Non-photosynthetic Orobanche Clade Sister to All Other Parasitic Orobanchaceae. Front Plant Sci. 2019;10. doi:10.3389/fpls.2019.00902

8.	McNeal JR, Bennett JR, Wolfe AD, Mathews S. Phylogeny and origins of holoparasitism in Orobanchaceae. American Journal of Botany. 2013;100: 971–983.

9.	Xia Z, Wen J, Gao Z. Does the Enigmatic Wightia Belong to Paulowniaceae (Lamiales)? Front Plant Sci. 2019;10: 528. doi:10.3389/fpls.2019.00528

10.	Yu W-B, Randle CP, Lu L, Wang H, Yang J-B, dePamphilis CW, et al. The Hemiparasitic Plant Phtheirospermum (Orobanchaceae) Is Polyphyletic and Contains Cryptic Species in the Hengduan Mountains of Southwest China. Front Plant Sci. 2018;9. doi:10.3389/fpls.2018.00142

11.	Schneider AC, Colwell AEL, Schneeweiss GM, Baldwin BG. Cryptic host-specific diversity among western hemisphere broomrapes (Orobanche s.l., Orobanchaceae). Annals of Botany. 2016;118: 1101–1111. doi:10.1093/aob/mcw158

12.	Camacho C, Coulouris G, Avagyan V, Ma N, Papadopoulos J, Bealer K, et al. BLAST+: architecture and applications. BMC Bioinformatics. 2009;10: 421.

13.	Katoh K, Standley DM. MAFFT multiple sequence alignment software version 7: improvements in performance and usability. Mol Biol Evol. 2013;30: 772–780.

14.	Minh BQ, Schmidt HA, Chernomor O, Schrempf D, Woodhams MD, Von Haeseler A, et al. IQ-TREE 2: New models and efficient methods for phylogenetic inference in the genomic era. Mol Biol Evol. 2020;37: 1530–1534.
